# Mixed Models Quantify Annual Volume Change; Linear Regression Determines Thalamic Volume as the Best Subcortical Structure Volume Predictor in Alzheimer’s Disease and Aging

**DOI:** 10.1101/2022.10.29.514239

**Authors:** Charles S. Leger, Monique Herbert, W. Dale Stevens, Joseph F. DeSouza, Alzheimer’s Disease Neuroimaging Initiative

## Abstract

**Background:** Thalamus-hippocampus-putamen and thalamus-cerebellar interconnections are dense. The extent this connectivity is paralleled by each structure’s volume impact on another is unquantified in Alzheimer’s disease (AD). Mixed model quantification of annual volume change in AD is scarce and absent inclusive of the cerebellum, hippocampus, putamen and lateral ventricles and thalamus. Among these structures, autopsy evidence of early-stage AD seems largely but not entirely restricted to the hippocampus and thalamus.

**Objective:** Variation in annual volume related to time and baseline age was assessed for the hippocampus, putamen, cerebellum, lateral ventricles and thalamus. Which subcortical structure’s volume had the largest explanatory effect of volume variation in other subcortical structures was also determined.

**Method:** The intraclass correlation coefficient was used to assess test-retest reliability of structure automated segmentation. Linear regression (*N* = 45) determined which structure’s volume most impacted volume of other structures. Finally, mixed models (*N* = 36; 108 data points) quantified annual structure volume change from baseline to 24-months.

**Results:** High test-retest reliability was indicated by a mean ICC score of .989 (*SD* = .012). Thalamic volume consistently had the greatest explanatory effect of hippocampal, putamen, cerebellar and lateral ventricular volume. The group variable proxy for AD significantly contributed to the best-fitting hippocampal linear regression model, hippocampal and thalamic longitudinal mixed models, and approached significance in the longitudinal lateral ventricular mixed model. Mixed models determined time (1 year) had a negative effect on hippocampal, cerebellar and thalamic volume, no effect on putamen volume, and a positive effect on lateral ventricular volume. Baseline age had a negative effect on hippocampal and thalamic volume, no effect on cerebellar or putamen volume and a positive effect on lateral ventricular volume.

**Interpretation:** Linear regression determined thalamic volume as a virtual centralized index of hippocampal, cerebellar, putamen, and lateral ventricular volume. Relative to linear regression, longitudinal mixed models had greater sensitivity to detect contribution of early AD, or potential AD pathology (MCI), via the group variable not just to volume reduction in the hippocampus but also in the thalamus.

## Introduction

Hippocampal atrophy, well documented in Alzheimer’s Disease (AD) (1, 2), is a morphological indicator associated with accelerated volume reduction in early AD (^3–6^).

Hippocampal atrophy is, as might be expected, strongly and positively correlated with histologically determined (hippocampal) neuron loss (7, 8). Where and when AD pathology begins remains elusive. Although hippocampal neuron loss and shrinkage undoubtedly contribute to amnestic dementia in AD (9, 10), AD involves neurodegeneration well beyond the borders of the hippocampus and other medial temporal lobe structures (^11, 12^). Moreover, accumulating evidence, outlined below, strongly suggests AD pathology research should, in addition to probing status of hippocampal and associated medial temporal lobe structures, include assessment of thalamic status in particular. Note, unless otherwise stipulated (i.e., histological, autopsy data stipulated), volume referred to in the present work is an imaging-based (mainly MRI) measure.

Braak and Braak’s seminal autopsy and staging (I-VI progressive stages) of AD revealed early evidence (stages II-III) of neuropathology in the form of abnormal tau formations (neuritic threads, plaques and neurofibrillary tangles) particularly in the anterior thalamic nuclei, and aspects of the entorhinal cortex and hippocampus with lesser such evidence in the striatum and cerebellum (^13, 14^). In A-C staging of extracellular amyloid accumulation, substantial amyloid deposits were revealed at stage C in several thalamic nuclei (anterior, reuniens, etc.,) but the hippocampus had, by comparison, much less neuropathology involvement (^14^).

Thalamic volume is reduced in the potential AD prodrome mild cognitive impairment (MCI) (^15, 16^). Thalamic volume is reduced in AD (^12, 17, 18^), particularly within the medial thalamic nuclei (which includes the anterior, mediodorsal, and reuniens nuclei) (^19^). In normal aging, structural alteration of the anterior thalamus is also indicated (^20^). Annualized volume reduction of the thalamus has been demonstrated second in extent only to annualized hippocampal volume reduction (4). Longitudinally, hippocampal atrophy progression/acceleration over 6-12 months occurs in AD but also in MCI (6). Hippocampal atrophy, particularly in AD but also in normal aging, is coupled with ventricular enlargement (^12, 21^).

There are inconsistent findings for altered putamen volume in AD. Putamen volume has been reported as unaltered in AD (^22^) as well as reduced in AD (4, 17). The cerebellum, implicated in cognitive AD dysfunction (^23^), has also demonstrated mixed volumetric findings in AD.

Cerebellar atrophy seems largely spared in AD(^24^) but has been reported as both greater than in normal aging (^25, 26^), and similar to that in normal aging in cross-sectional and longitudinal analyses (^27^).

Volume of any structure is integrated with a physiological system that includes connectivity. Thalamic connectivity, particularly diverse, has bidirectional or looping circuits involving cortical regions, basal ganglia, and limbic (e.g., hippocampus, posterior cingulate cortex, fornix, piriform cortex, amygdala) structures (^11, 28^). There is also a plethora of reciprocal cerebellar-thalamic connections (^29–32^). Recent diffusion-weighted imaging, incorporating a large data sample (*N* = 730), has substantiated the elaborate and dense fiber pathways interconnecting the thalamic anterior nuclei and the hippocampus as well as other limbic system structures (^33^).

Moreover, as concluded in meta-analytic research involving over ten-thousand human brain imaging studies, focal lesions of the thalamus have widespread repercussions, disrupting diverse cortical functional networks, further exemplifying the hub-like role of the thalamus (^34^). In short, the thalamus is diversely interconnected with other structures; dysfunction in the thalamus, or in one or more of its subdivisions, has demonstrated impact extending outside thalamic borders (^35, 36^).

The current work used normalized volume in early AD, potential AD (MCI) and controls (CN) of five structures: cerebellum, hippocampus, putamen, lateral ventricles and thalamus.

These structures, as reviewed, can be impacted by AD or normal aging. Separate linear and longitudinal mixed model regression analyses assessed volume of the structures. Notable, three of the structures (cerebellum, hippocampus, putamen), as reviewed, are densely interconnected with the thalamus, or with respect to the lateral ventricles anatomically juxtaposed to the thalamus. Further, early stage AD-type pathology (abnormal tau formations) is localized largely, though not entirely, to the hippocampus and thalamus (^13, 14^). In short, thalamic connectivity (^34^), thalamic early-stage AD involvement (^13, 14^) and volume change (^12, 18, 19, 37^) combine to suggest the potential of this single structure to inform on the status of the other four structures (cerebellum, hippocampus, putamen, lateral ventricles) in early AD. As already noted, (normalized) volume was the metric of interest. Volume served as the response variable (univariate) in five distinct linear and five distinct mixed models. Explanatory variables were selected from the literature and refined in regression procedures. Linear regression explanatory variables were initially baseline age, gender, cognitive measures, the group variable (AD, MCI, CN) as well as the volume of the other structures. The AD and MCI groups, as defined by the Alzheimer’s Disease Neuroimaging Initiative (^45^), represented early AD and potential AD (MCI); the group variable AD and MCI cohorts, served as a proxy for early AD or MCI. In linear regression models, each structure’s volume served as a response variable once and as an explanatory variable four times (given five structures other than the response variable). This, model-based approach, assessed contribution of a given predictor, including another structure’s volume, to predict/explain volume of the response variable structure volume. Mixed model explanatory variables were effects of time, baseline age and group. As with linear regression, each structure’s volume model determined explanatory variables contributing most to volume. Evidence of early stage hippocampus and thalamus (^13, 14^) AD-type pathology prompted the first avenue of inquiry: does the presence of early-stage hippocampal and thalamic AD pathology account for greater AD-related variation in hippocampal and thalamic volume compared to AD-related volume variation in the other three structures? Indication of early-stage AD pathology particularly in the thalamus and hippocampus (^13, 14^) coupled with the dense interconnections among these structures (^33, 38–40^) implies a possibly linked hippocampus-thalamus neuronal fate in early-stage AD. As such, significant explanatory effects/estimates of the group variable AD and MCI cohorts were expected for thalamic and hippocampal volume regression models (linear and mixed). In a second vein of inquiry, linear regression determined if any single structure’s volume as an explanatory variable accounted consistently for more variation in response structure volume. It was expected that thalamic volume as an explanatory variable would parallel the central and impactful effect of thalamic connectivity (briefly outlined in the preceding paragraph) in normal aging and early AD. To the best of our knowledge, study objectives and methods (see Materials and methods) are unique within the context of early AD investigation.

## 2. Materials and methods

### Procedures and power analyses

The main measure of interest was, the biological indicator, MRI-based subcortical structure volume of the hippocampus, cerebellum, putamen, hippocampus and thalamus.

Subcortical structure segmentation/volume estimates were completed using the dedicated volumetry platform volBrain (^41^) (https://volbrain.upv.es). As outlined in the preceding statement, regression models (univariate) were the principal method of assessment. Prior to regression analyses test-retest reliability of volBrain (^42^) subcortical segmentation was determined using the intraclass correlation coefficient (ICC). Prior to the mixed model analysis a repeated-measures correlation (^43^) analysis was conducted. Finally, subsequent to mixed model analyses, a simple annualized volume calculation was completed (see Supporting information II for details).

Linear and mixed model power analyses were conducted to estimate the minimum number of data instances required to provide at least 80% power to reject the null hypothesis with an appropriate effect size. An initial power analysis for the linear multiple regression models indicated a model with 2 to 3 predictors, a large .80 effect size and 80% power at α= .05 would require *N* = 45 subjects (at least 13 per group, where analyses included 3 groups).

Accordingly, the preference was to obtain a baseline study sample of at least 45, with 15 unique records per group (AD, CN, MCI). The actual power achieved in the linear model results ranged from a low .85% with an adjusted *R^2^* of .25 to a high for the hippocampus and thalamic models of .99% and an adjusted *R^2^* of between .52 and .67.

With respect to a longitudinal mixed model power analysis, some loss of records was expected across time. As such, we ran some preliminary mixed model simulations (^44^) with an *n* = 30 (unique records) across 3 time points (90 data instances), one fixed effect, time, and the subject intercept random effect. Simulations with a conservative effect size of -0.07 using the full 90 data instances had a power of 92.50%. The actual longitudinal data set proved to have a range of 33-36 unique complete cases of data (99-108 data instances over 3 time points), fixed effects estimates ranged from -.13 for the hippocampal model AD group dummy variable to nil for the putamen model. Actual mixed model pseudo *R^2^* (*pR^2^m= fixed effects*, *pR^2^c = fixed plus random effects*) ranged from *pR^2^m* .53 and *pR2c* .99 in the hippocampal model to nil *pR^2^m* and *pR^2^c* .97 in the putamen model.

### 2.2 Subject data

Data was sourced from the Alzheimer’s Disease Neuroimaging Initiative (^45^) (ADNI) database. Specifically, ANDI 1 and ADNI GO data was used. ADNI is a public-private partnership initiated in 2003, and led by principal investigator Michal W. Weiner MD. The main objective of ADNI has been to investigate whether serial magnetic resonance imaging (MRI), positron emission tomography (PET), other markers (biologics, clinical and neuropsychological assessments) can be combined to quantify progression of early Alzheimer’s disease (AD) and mild cognitive impairment (MCI). Up to-date information can be found at www.adni-info.org.

ADNI’s unique back-to-back (BTB) baseline and 12-month same-subject scan pairs were used to assess subject intra-session scan segmentation test-retest reliability (see 2.5). These BTB scans are same-subject T1 images taken minutes apart resulting in baseline and month twelve pairs of scans per subject. Using ADNI’s online image Advanced Search interface, the following criteria were used: original, pre-processed and processed data; ADNI 1, ADNIGO; AD, CN MCI cohorts, present at baseline and 12 months; age range 55-88; MMSE and CDR included; 3T; T1. This returned 3311 scans of which 156 scans represented distinct, individual subject data. During inspection of scans, artifacts were most common in those subjects who had multiple repeats of scans at baseline and at 12-month time points. After filtering-out those subjects with greater than 10 scans per time point, the pool of subjects was reduced to 72. After a final and closer inspection of these scans, 47 were selected. Of these, volBrain processing errors occurred for two MCI cohort members (023_S_1126 and 023_S_1247). This left an intra-session dataset for test-retest reliability (see section 2.5) involving 180 scans from 45 subjects (15 per AD, CN and MCI groups times 4 BTB intra-session scan pairs).

Linear regression analyses (see section 2.5) used baseline scans and data from 45 subjects: 15 AD, 12 female; 15 healthy age-matched CN, 7 female; and 15 scans from the MCI group, 5 female. The AD cohort was relatively early stage, pre-dementia or mild dementia based on ADNI classification.

With regard to the longitudinal mixed model dataset, nine cases present in the original baseline (*n* = 45) and 12-month data were absent in the 24-month data. This resulted in a final longitudinal 36-subject dataset with complete case data at all three time points: 10 AD, 14 CN, and 12 MCI at each of the baseline, 12-month and 24-month timepoints. It is acknowledged mixed model intercepts could have benefited by utilizing all 45 cases with available data at baseline and 12-months. Nevertheless, in this analysis, only complete cases, the 36 cases present at all time points, were to be used in the longitudinal mixed model analyses. Further, from this group of 36 complete cases, the final longitudinal data set had a final range of 33-36 unique complete cases of data (99-108 data instances over 3 time points). With the exception of the lateral ventricular mixed model, 1 to 3 data instances in mixed models proved overly influential. These cases were removed. See *Supporting information VI* for Mixed model diagnostics details.

Clinical and demographic data were utilized in regression analyses that followed volBrain intra-session segmentation test-retest reliability assessment. Cognitive measures were the Mini-Mental State Examination (MMSE: maximum score of 30 higher is better) (^46^), and the Clinical Dementia Rating (^47, 48^) scale (CDR: normal = 0, and .5, 1, 2, and 3 respectively reflect very mild, mild, moderate and severe dementia). The demographic variables were age and gender. However, notable, among the explanatory variables included in models initially, gender did not make a significant contribution to any model. In addition, there was a large imbalance in gender, especially for the AD group. To avoid misleading regression estimates gender was not included in models.

### 2.3 MRI acquisition

Acquisition of ADNI MRPAGE pulse sequences was highly standardized (^49^) across multiple sites utilizing GE, Philips or Siemens equipment: baseline, month-12, and 24-month same-subject scans employed the same sequences at 3T. MPRAGE T1 preprocessing was restricted to conversion from DICOM to NIFTI format. The slice thickness for these samples was consistently 1.2 mm and all pixels were square. Voxel volume range was 1.20 mm^3^ to 1.24 mm^3^. All images were carefully inspected for artefacts.

### 2.4 Automated segmentation

VolBrain (^42^) is an online platform (http://volbrain.upv.es) dedicated to volumetric analyses. A so-called patch-based method, it has a fully automated pipeline based on fusion of multiple atlases (^41, 42^). We used the default volBrain library, which consists of 50 T1-weighted images (MPRAGE and SPGR). The volBrain pipeline consists of several steps: denoising, inhomogeneity correction, registration of affine to MNI space, fine inhomogeneity correction, intensity normalization, intracranial cavity extraction, tissue classification, non-local hemisphere segmentation, and subcortical non-local segmentation.

### 2.5 Statistical analyses

As outlined in section 2.2, two baseline (V0_A_ and V0_B_) and two 12-month (V12_A_ and V12_B_) same-subject scan pairs were used to assess test-retest reliability of these intra-session scans. Segmentation test-retest reliability was assessed with the intraclass correlation coefficient (ICC) for agreement using same-subject (intra-session) baseline and 12-month scan pairs. ICC was used to assess agreement between baseline (same subject) scan pairs V0_A_ and V0_B_ and then between V12_A_ and V12_B_ scan pairs. The ICC was calculated by group for both left and right hemisphere using raw structure volume.

Regression analysis were conducted using normalized volume of the hippocampus, putamen, thalamus, lateral ventricles and the cerebellum. Normalized volume was a relative measure: a ratio of structure total (left plus the right) volume to total intracranial volume (ICV). Normalized volume values are a percent: e.g., structure volume/ICV * 100. The type I error rate for regression analyses was set at .05 (α = .05). As always, regression coefficients quantify the magnitude or strength of the predictor/explanatory/independent variable effect on outcome. The extent of any explanatory variable effect, the extent that it explains variation in outcome, should not be confused with mediation in this research – this research uses regression not mediation.

#### 2.5.1 Linear regression methods

Each structure’s volume served as a response variable once and as an explanatory variable four times (given five structures). Having each structure serve once as the response variable but otherwise in an explanatory variable role allowed for structure/model-based determination of which structure’s volume most effected volume of other structures. Along with volumes of structures, linear regression explanatory variables were baseline age, cognitive measures

(MMSE, CDR) and the group variable, where AD and MCI cohorts, served as a proxy for early AD or MCI. Owing to the gross imbalance in cell size with respect to gender (e.g., the baseline AD group had 12 females but just 3 males; and the 24-month data AD data had 9 females but just one male, see *2.1*), and to avoid a misleading outcome, gender was not used as a covariate. Stepwise regression (backward, using AIC) was used to aid in selection optimal final model features. All final linear regression models complied with regression assumptions and there was an absence of overly influential observations. It warrants note however that the group and clinical dementia rating (CDR) explanatory variables were too highly correlated (*rs* = .87) to include in the same model, hence these two variables were examined in two distinct regression analyses. Only models with hippocampal volume as the response variable retained the CDR and group variables. Two linear regression hippocampal models were conducted: one using the group variable and one using CDR. For each linear model, 2-way interactions were assessed for explanatory volume variables by age and explanatory volume variables by group in final models retaining the variables. It warrants note that each of the five structures has as a differing volume range or scale of values, making direct comparison among structure RMSE values inappropriate. To aid in comparison, a normalized version of RMSE, NRMSE, was also calculated: the NRMSE was approximated by dividing the RMSE by the standard deviation of Y. Standardized estimates/coefficients, also provided, are unitless and therefore facilitate comparisons of differently scaled variables(^50^). The principal of parsimony was adhered to for both linear regression and mixed linear regression models: if the addition of an explanatory variable did not significantly improve the fit of the model to the data, the term was dropped and the simpler model retained.

#### 2.5.2 Linear mixed model methods

In mixed longitudinal models, the response variable was, as in linear regression, the volume of a single structure. Mixed models focused on quantifying explanatory variable contribution (notably the group variable AD and MCI cohorts that served as proxies for early AD or MCI.) to annual volume change in the same five structures over two years. The mixed model explanatory variables were effects of time, baseline age and group. Prior to conducting mixed model analysis, a repeated-measures correlation analysis was conducted to assess the nature of bivariate relationships between variables. The software package used (^43^) calculates a repeated-measures coefficient (*r*_rm_) that has a -1 to 1 index of association strength between two variables analogous to Pearson *r*, but unlike the latter does not violate the assumption of observation independence; non-independence is accounted for. The *r*_rm_ provides quick longitudinal insight. Compared to a mixed model, however, there are two main limitations. First, as with Pearson correlation, only the relationship between two variables can be assessed simultaneously. Second, it is analogous to the mixed null model, including only the random (varying) subject intercept (mean) and the overall slope (intercept) (^43^).

The linear mixed algorithm models account for dependence (non-independence of residuals) by including random, individual subject idiosyncratic variability in mean volume (the intercept) as well as subject variability in the effect of time on volume. The mixed longitudinal volume analyses, as with the linear regression analyses, involved separate analysis for hippocampal, putamen, thalamus, cerebellum and lateral ventricular volume. Also similar to the linear model procedures, the mixed model response variable was normalized volume of one of the latter structures. However, unlike the procedure for the linear models, volumes of the structures other than the response variable did not serve as explanatory variables. The mixed model fixed effect explanatory variables were time (baseline, 12-month, and 24-month time points) baseline age and group. Time was included as a numeric variable, an approach adopted in a much-cited paper (^51^). Age and time points will covary when time and age are included in a longitudinal analysis. To control time-age collinearity, static baseline age was used. Baseline age and time (for all 3 time points) were centered to assess interactions. A time by group (AD, CN, MCI) interaction was examined for each model. Random effects consisted of individual subject intercept and the individual subject slope of time.

As in the linear regression models, in the mixed model analysis volume of one of five subcortical structures (the hippocampus, cerebellum, putamen, lateral ventricles or thalamus) served as the response variable in separate models. Mixed model independent (fixed effect) main effect variables consisted of baseline age, time and group.

Separate mixed models were executed in the recommended sequence (^52, 53^). Specifically, mixed models began with the simplest model and progressed in complexity. Seven different levels of complexity (i.e., model types 1 to 7, see below), were executed using the total normalized volume of each structure as the response variable. Only the final model with the best, most parsimonious fit to the data was reported in detail. However, a comparison of all model type results is provided in *Supporting information I*. Again, for each of the five structures, seven different model types were executed with 1-7 levels of complexity. Initially, an intercept only (unconditional) model was assessed, a model where the intercept did not vary (volume was simply regressed on the constant 1). This unconditional model was executed using generalized least squares (model 1). Next, another unconditional model was executed in which intercepts were random, permitting means to vary across subjects (model 2). This adds to the overall model intercept the effect of subject-specific intercepts. This accounts for structure volume nested within subjects (individual variation in volume). In model 3, time was added as a random effect of subject. The random slope of time for each subject assessed the extent the effect of time on volume differs for each subject. Again, bear in mind, that longitudinally, the effect of time is nested within subjects. Next, in model 4, time was added as a fixed effect, which as in standard linear regression, estimates, on average, the effect of time on volume across all individuals. The next level of model complexity, model 5, added the fixed effect of baseline age. The model 6 added the effect of group (AD, CN, MCI) to determine the effect of group membership on volume; AD vs controls (the reference group) and MCI vs controls. Finally, a model 7 was added that involved a group by time interaction. In addition, a quadratic fixed effect time variable was also included to assess potential quadratic, accelerated, effect of time (i.e., time^2^) in the model.

Adding a random quadratic term for time to the random effects of time and subject is optional; a quadratic effect of time can be included as a fixed only or as both a fixed and random effect (^52^).

In general, whether it necessary to include all fixed effects also as random effects is a subject of debate that can be reviewed various statistics forums.

Measures of model fit, for a given structure’s longitudinal volume assessment, used to compare across model types (1to 7 as described above), were Akaike Information criterion (AIC), deviance (or loglikelihood) and chi-squared statistic (or a likelihood ratio test), RMSE and NRMSE and pseudo *R^2^* (^54^). The latter pseudo *R*^2^ has two components: *pR^2^*m, which refers to the fixed effects (marginal) contribution and *pR^2^*c, which refers to total fixed plus the random effects contribution to pseudo variance accounted for by the model. The chi-squared statistic was used for model type (1 to 7) comparison. To facilitate comparison of models the maximum likelihood (ML) method of estimating mixed model parameters was used, though it is recognized that restricted maximum likelihood (REML) has demonstrated more precise and less biased random effect variance component estimates (^55^). Also, formal pairwise tests, where appropriate, were conducted on final models using a Tukey test. In addition, a standard Cohen’s *D* effect size was adopted to assess pairwise test effect size. The standard Cohen’s D ignores design (between-subject, mixed model etc.) information. While mixed-model specific effect sizes methods can be adopted it has been convincingly argued that such tests impair comparison of effects across differing analysis designs (^56^).

#### 2.5.3 Regression model diagnostics

To mitigate against overfitting, all linear and mixed regression models were compliant with a relaxed one predictor to five event (data instance) rule of thumb (^57^). Four of the linear regression models complied with the more stringent one predictor to ten events rule of thumb (^58^), and three of five mixed models complied with one predictor to ten events rule of thumb (see *Supporting information VI* for mixed model diagnostic details). All model residuals were assessed for regression assumption violation (including mixed model random effects) and the presence of influential cases. Note, two mixed model packages were used, package lme4 (^51^); package nlme (^59^). This served as a check on analyses and also offered some methodological latitude. Analyses were conducted in *R* (^60^). on a 4-core, (3.2 GHz, 6 GB) Apple OS Higher Sierra (v. 10.13.6) system.

## 3.0 Results

FIG 1 (A) depicts the subcortical structures of interest in MNI 152 space. FIG 1 (B) displays the color coding for structures. Notable, only volumetry of whole structures was used. However, FIG 1 (A) also localizes thalamic anterior nuclei (red segments on the anterior/superior aspect of the turquoise-colored thalamus in FIG 1 A). The anterior thalamic nuclei group are localized in FIG 1 (A) because they were referred to in the introduction and will be included in the next phase of this project. VolBrain parcellation is provided in FIG 1 (C). The volBrain parcellation in FIG 1 (C) includes, by default, other structures not assessed in the current work. Only structures matching the color coding in (B) are relevant.

**FIG 1.**
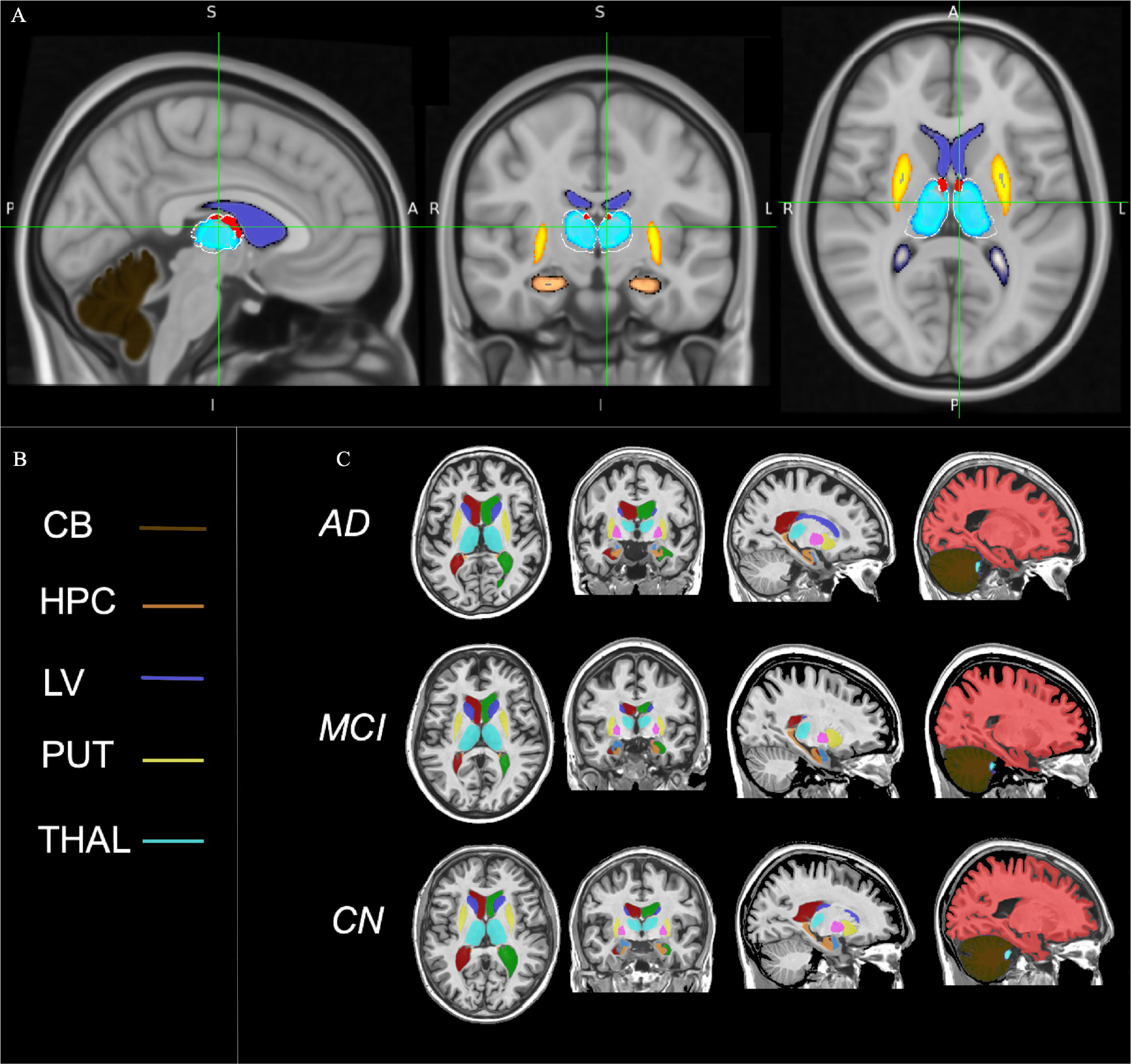
Subcortical structure mapping in MNI 152 space (.5mm). (A) Three cardinal planes MNI -3.93, -13.61, 9.29; x=94, y = 112, z= 81; (B) color coded legend; (C) VolBrain Subcortical parcellation in MNI 152 space. AD = Alzheimer’s disease; MCI = mild cognitive impairment; CN *= controls CB = cerebellum (dark brown); HPC=hippocampus (Copper); PUT = putamen(yellow); LV = lateral ventricles (purple); THAL = thalamus (turquoise; surrounded by white boarder in A). Structures in A derived from Harvard-Oxford Subcortical Structural Atlas. Anterior thalamic nuclei group (red) and the lateral dorsal thalamic nucleus (red) (Krauth et al., 2010). Note, the lateral dorsal nucleus is often included as part of the anterior thalamic nuclei group. Radiological convention using FSLeyes version 1.3.0*.

**Table 1** contains baseline demographic, MMSE, CDR, as well as raw, non-normalized, volume measures in mm^3^. Measures are arranged by group (AD, MCI, CN). The volBrain volume estimates are consistent with other AD research (^61^) where cohorts were of similar age as those in the current work.

**TABLE 1:** Demographics, MMSE, CDR, baseline volume, N=45

| | AD ($n = 15$) | CN ($n = 15$) | MCI ($n = 15$) |
| --- | --- | --- | --- |
| *Measures* | *Mean (SD)* | *Mean (SD)* | *Mean (SD)* |
| Age | 74 (8) | 74 (8) | 73 (8) |
| CDR (median, range) | .5 (.5, 1) | 0 (0, .5) | .5 (.5, .5) |
| MMSE | 22 (5) | 29 (1) | 25 (3) |
| *Baseline volume mm<sup>3</sup>* | *Mean (SD)* | *Mean (SD)* | *Mean (SD)* |
| CB | 59267 (10454) | 61901 (4202) | 64346 (9631) |
| HIPP | 2942 (635) | 3783 (485) | 3216 (549) |
| LV | 20532 (16096) | 14179 (7031) | 23840 (12776) |
| PUT | 3971 (669) | 3980 (413) | 3914 (508) |
| THAL | 4811 (620) | 5094 (590) | 5025 (472) |

AD = Alzheimer's disease; MCI = mild cognitive impairment; CN = controls; CDR = cognitive dementia rating: 0 – 5, 0 is normal; MMSE = Mini-Mental State Examination: maximum of 30 points, 25–30 normal cognition, 21–24 mild dementia, 10–20 moderate dementia, $\leq 9$ severe dementia; THAL = thalamus; HIPP = hippocampus; PUT = putamen; volumes are baseline structure averages across hemispheres in mm<sup>3</sup>, e.g., left CB + right CB/ 2; values are mean and SD unless otherwise indicated. All volume measures in this table are in native space.

Intraclass correlation coefficient (ICC), test-retest reliability was high as indicated by the mean ICC score of .989 (*SD* = .012). The lowest ICC was .937 (95% CI .829, .978) for the 12- month time-point measure of the left hippocampal hemisphere in controls. For details on all ICC measures see Table S1-11 in *Supporting Information I*.

Group measures of baseline age, gender MMSE, and CDR were initially assessed.

Binomial tests within AD, controls and MCI groups indicated AD cohort gender imbalance; a proportion of .80 females (12/15 = .80 female), which significantly differed from the expected proportion of .5 (50%), *p* = .035 (95% CI .52, .96). Because of this gender imbalance, noted earlier, and its potential to bias models, gender was not included as a variable. ANOVA indicated age (which did not violate homogeneity of variance across groups) did not significantly differ among groups, *F*(2, 42) = .87, *p* = .26. Kruskal-Wallis tests were used to test MMSE and CDR. Each of the latter two variables had non-normal distributions and high heteroscedasticity. MMSE scores did significantly differ among groups, χ^2^ (2) = 26.9, *p* < .0001, *r^2^ ^adj^* = .59 (*r^2^ ^adj^* ANOVA on ranks). A posthoc Dunn Test, controlling for multiple comparisons (Benjamin-Hochberg), indicated AD (*p* < .0001) and MCI (*p* < .001) groups significantly differed from controls. The MMSE mean scores of AD and MCI cohorts did not significantly differ, *p* = .30. The pattern was the same for CDR scores. CDR mean scores significantly differed among groups, χ^2^ (2) = 35.2, *p* < .0001, *r^2^ ^adj^* = .79. A post hoc Dunn’s Test, controlling for multiple comparisons (Benjamin-Hochberg), indicated AD and MCI groups significantly differed from controls (*p* < .0001). The CDR mean scores of AD and MCI cohorts did not significantly differ, *p* = .09.

### 3.1 Normalized volume by group

All structure volume measures used in the regression analyses were normalized, where normalized refers to a given structure’s volume relative to total intracranial volume (ICV); values are percentages – i.e., structure volume/ICV * 100. **Table 2** specifies volBrain normalized structure volume for the baseline dataset used in the linear regression analyses (*N*= 45: AD= 15, CN=15, MCI =15) and for the longitudinal 36-subject dataset (AD = 10, CN = 14, MC =12) used in mixed model analyses. The longitudinal 36-subject dataset had complete data instances across baseline, 12-month, and 24-month time-points. Again, structure volume is provided as a percentage of total intracranial volume. The median longitudinal volume values tabulated in **Table 2** convey apparent small reductions in hippocampal and thalamic volume across groups, with less discernable 24-month change in putamen and cerebellar volume. Also indicated in the longitudinal data, the lateral ventricles had a pattern of increasing size over 24 months.

**TABLE 2:** Normalized structure volume as a percentage of total intracranial volume; median and range

|  | *HPC* | *PUT* | *THAL* | *LAT VENT* | *CB* | *AGE* | *CDR* | *MMSE* |
| --- | --- | --- | --- | --- | --- | --- | --- | --- |
|  | *MDN*<br>*(rng)* | *MDN*<br>*(rng)* | *MDN*<br>*(rng)* | *MDN*<br>*(rng)* | *MDN*<br>*(rng)* | *MDN (rng)* | *MDN (rng)* | *MDN (rng)* |
| **Gp Time** |  |  |  |  |  |  |  |  |
| **<u>Baseline data, N = 45: 15 AD, 15 CN, 15 MCI</u>** |  |  |  |  |  |  |  |  |
| AD base | **.42**<br>(.29, .58) | **.59**<br>(.50, .70) | **.72**<br>(.56, .84) | **2.42**<br>(.96, 8) | **8.86**<br>(6.67, 10) | **72**<br>(57,86) | **.5**<br>(.5, 1) | **24**<br>(13, 29) |
| CN base | **.56**<br>(.41, .67) | **.58**<br>(.51, .64) | **.79**<br>(.60, .84) | **1.68**<br>(.68, 4) | **8.96**<br>(7.74, 10.6) | **73**<br>(60, 86) | **0**<br>(0, .5) | **30**<br>(27, 30) |
| MCI base | **.43**<br>(.34, .59) | **.55**<br>(.41, .63) | **.69**<br>(.59, .81) | **3.05**<br>(1, 5.5) | **8.63**<br>(7.02, 11) | **73**<br>(56,88) | **.5**<br>(.5, .5) | **26**<br>(19, 29) |
| **<u>Longitudinal (3 time-points): AD n = 30 (10 x 3), CN n = 42 (14 x 3), MCI n = 36 (12 x 3)</u>** |  |  |  |  |  |  |  |  |
| AD base | **.41**<br>(.29, .57) | **.59**<br>(.50, .69) | **.70**<br>(.56, .82) | **2.7**<br>7(.96, 5.6) | **8.91**<br>(6.7, 10) | **72**<br>(64, 85) | **1**<br>(.5, 2) | **24**<br>(13, 29) |
| AD m12 | **.38**<br>(.27, .57) | **.58**<br>(.49, .69) | **.70**<br>(.58, .81) | **3.25**<br>(.87, 6.13) | **8.8**<br>(6.8,9.9) | **73**<br>(65, 86) | **1**<br>(.5, 2) | **24**<br>(13, 29) |
| AD m24 | **.36**<br>(.25, .56) | **.56**<br>(.47, .68) | **.68**<br>(.56, .80) | **3.73**<br>(1.2, 7.93) | **8.7**<br>(6.7, 9.9) | **74**<br>(66, 87) | **1**<br>(.5, 2) | **22**<br>(10, 27) |
| CN base | **.55**<br>(.41, .62) | **.58**<br>(.51, .64) | **.79**<br>(.60, .84) | **1.67**<br>(.68, 3.72) | **8.9**<br>(8, 10) | **73**<br>(60, 86) | **0**<br>(0, .5) | **30**<br>(27, 30) |
| CN m12 | **.55**<br>(.42, .62) | **.57**<br>(.50, .64) | **.75**<br>(.61, .82) | **1.98**<br>(.70, 3.73) | **8.8**<br>(7.9, 10.5) | **74**<br>(61, 87) | **0**<br>(0, .5) | **30**<br>(27,30) |
| CN m24 | **.52**<br>(.41, .61) | **.57**<br>(.52, .66) | **.72**<br>(.59, .83) | **2.28**<br>(.75, 3.98) | **8.9**<br>(7.8, 10.6) | **75**<br>(62, 88) | **0**<br>(0, .5) | **30**<br>(27, 30) |
| MCI base | **.45**<br>(.34, .59) | **.56**<br>(.47, .63) | **.69**<br>(.59, .81) | **2.56**<br>(1.0, 5.51) | **8.88**<br>(7.02, 10.8) | **72**<br>(56, 83) | **.5**<br>(.5, .5) | **26**<br>(19, 29) |
| MCI m12 | **.44**<br>(.35, .57) | **.56**<br>(.46, .65) | **.68**<br>(.58, .81) | **2.96**<br>(1.21, 5.61) | **8.91**<br>(7.2, 10.7) | **73**<br>(57, 84) | **.5**<br>(.5, .5) | **26**<br>(19, 29) |
| MCI m24 | **.44**<br>(.34, .57) | **.56**<br>(.46, .67) | **.66**<br>(.59, .81) | **3.19**<br>(1.31, 6.33) | **8.9**<br>(7.4, 10.5) | **74**<br>(58, 85) | **.5**<br>(.5, 1) | **25**<br>(11,30) |

AD = Alzheimer's disease group; CN = controls; MCI = mild cognitively impaired CB = cerebellum; HPC= hippocampus; PUT = putamen; LAT VENT = lateral ventricles Gp = group; base = baseline; m12 = 12 months post baseline; m24 = 24 months post baseline; MDN = median; rng= range; Structure volume is a percentage of total intracranial normalized volume: e.g., for the median AD longitudinal baseline hippocampus volume, left + right hippocampal volume / total intracranial volume \*100 = .41.

FIG 2 provides additional insight into volume change between time points (baseline vs. 12-months; baseline vs 24-months; 12-months vs. 24-months) within groups. Structure volumes in FIG 2 reflect normalized volume (e.g., left + right hippocampal volume / total intracranial volume *100), Normalized structure volume was converted to a z-score to facilitate across structure comparison. FIG 2 panels reflect hippocampal (A), lateral ventricular (B) and thalamic (C) volume between time point tests (baseline, 12-months and 24-months) for each group. False discovery rate was used to adjust p-value values for multiple comparisons. Plot values (x-axes) are in standard deviations. The same volume pairwise tests between time points for cerebellar and putamen volume is provided in Table S1-24 (see Supporting information I). Table S1-24 will indicate significant (*p* < .05) putamen volume changes only for the AD cohort between each time point but no significant cerebellar volume differences between time points for all groups.

**FIG 2.**
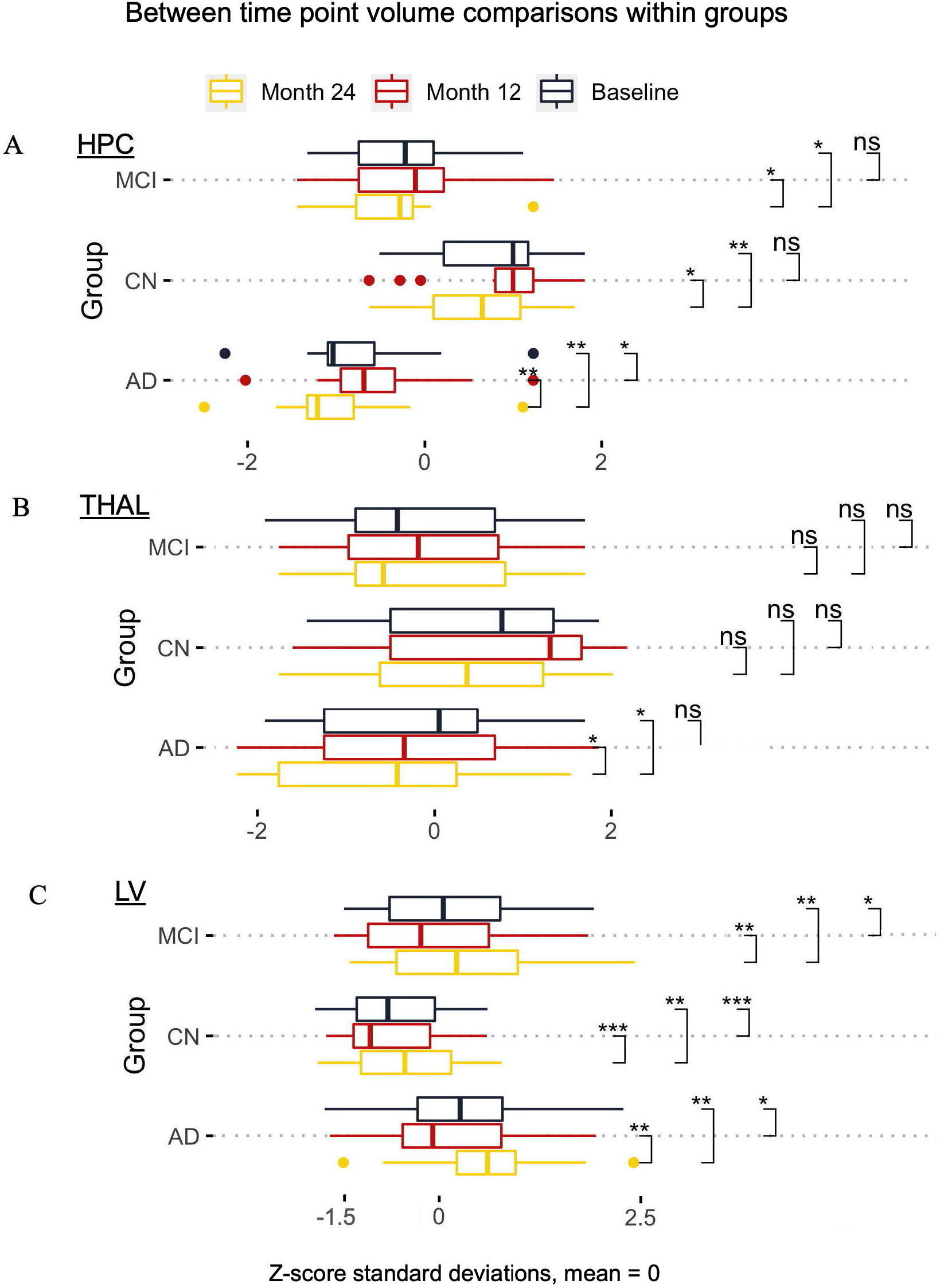
Between time-point comparisons. AD = Alzheimer’s, CN = controls, MCI = mild cognitively impaired. X-axes in panels X-axes reflect the standard deviation (mean = 0) of normalized volume (e.g., left + right hemisphere volume / total intracranial volume *100); y-axis indicates group; time is color coded. Panels A, B and C are the respective hippocampal (HPC), thalamus (THAL) and lateral ventricular (LV) normalized volumes as z-scores; *= p < .05; ** = p < .01; *** = p < .001. Note, the non-normality of lateral ventricular volume required robust (Wilcox) pairwise tests. Plots based on longitudinal baseline, 12-month and 24-month data.

### 4.1 Linear regression volume analyses

Linear regression analyses were conducted using baseline data. This dataset had 45 subjects, 15 in each group. See section *2.5.1* *Linear regression methods* for more method information (additional details are available in *Supporting Information IV*). The main objective was to determine the combination of variables (structure volume, age, group, and MMSE) that best explained a given structure’s volume (see section *2.5.1* *Linear regression methods*). Results of all the final (best fitting) models) are detailed in **Table 3**. Tabulated values reflect the effect of a given predictor on the outcome measure (fractional measure: structure volume / total intracranial volume * 100). Only the models mitigating against overfitting with the best data fit (based on criteria including model RMSE, *adjusted-R**^2^,*** and variable estimates) are reported in **Table 3**. The thalamus (normalized volume), as an explanatory variable (i.e., that explains/predicts outcome), had the largest significant estimate in all models. Model predicted means were as follows: cerebellar volume 8.83% (95% CI 8.61, 9.05); putamen volume, .573% (95% CI .56, .59); lateral ventricles, 2.35% (95% CI 2.12, 2.60); thalamus, .727, (95% CI .71, .74). A range of model volume predicted values (e.g., using first and third quarter predictor data values) is available on request.

**TABLE 3:**
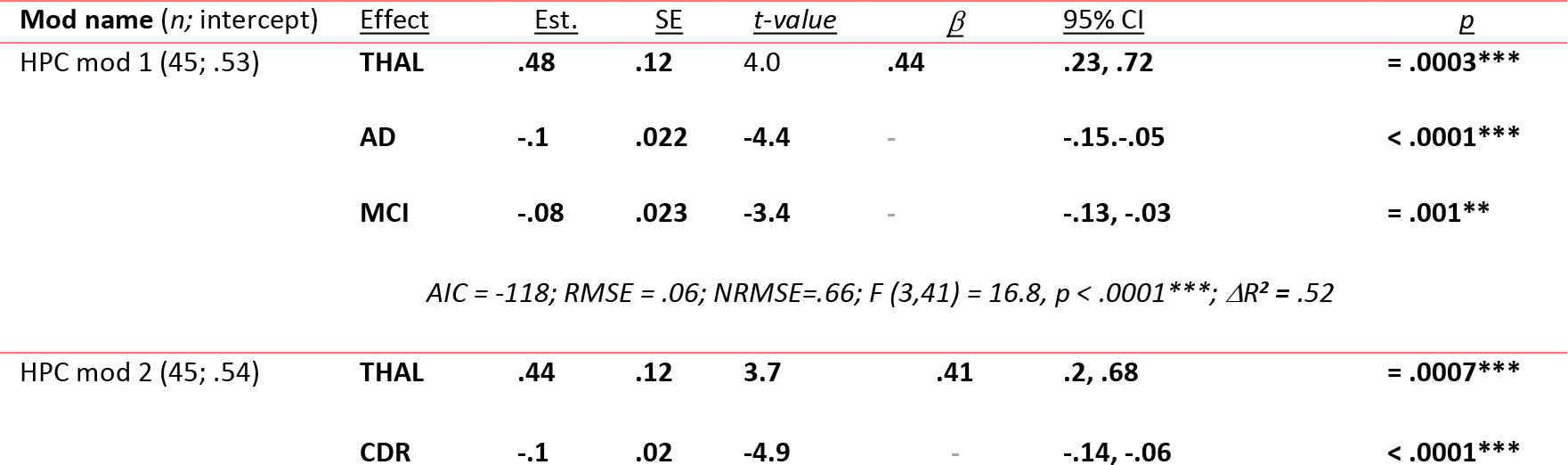
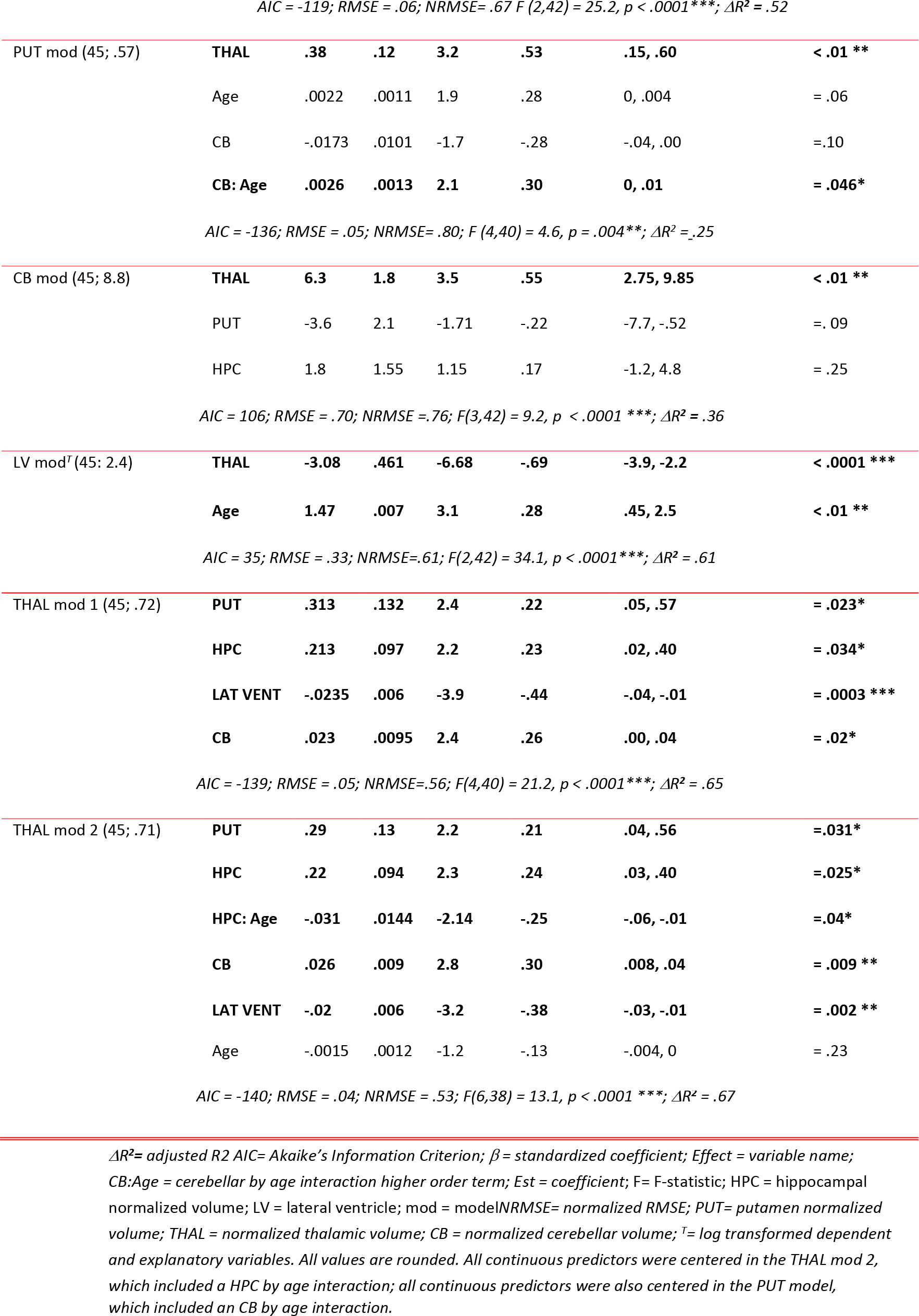
Linear regression model results

| Mod name (n; intercept) | Effect | Est. | SE | t-value | $\beta$ | 95% CI | p |
| --- | --- | --- | --- | --- | --- | --- | --- |
| HPC mod 1 (45; .53) | THAL | .48 | .12 | 4.0 | .44 | .23, .72 | = .0003*** |
|  | AD | -.1 | .022 | -4.4 | - | -.15, -.05 | < .0001*** |
|  | MCI | -.08 | .023 | -3.4 | - | -.13, -.03 | = .001** |
| AIC = -118; RMSE = .06; NRMSE=.66; F (3,41) = 16.8, p < .0001***; $\Delta R^2$ = .52 | | | | | | | |
| HPC mod 2 (45; .54) | THAL | .44 | .12 | 3.7 | .41 | .2, .68 | = .0007*** |
|  | CDR | -.1 | .02 | -4.9 | - | -.14, -.06 | < .0001*** |
| $AIC = -119$ ; $RMSE = .06$ ; $NRMSE = .67$ $F(2,42) = 25.2$ , $p < .0001^{***}$ ; $\Delta R^2 = .52$ |  |  |  |  |  |  |  |
| PUT mod (45; .57) | **THAL** | **.38** | **.12** | **3.2** | **.53** | **.15, .60** | **< .01 \*\*** |
|  | Age | .0022 | .0011 | 1.9 | .28 | 0, .004 | = .06 |
|  | CB | -.0173 | .0101 | -1.7 | -.28 | -.04, .00 | = .10 |
|  | **CB: Age** | **.0026** | **.0013** | **2.1** | **.30** | **0, .01** | **= .046\*** |
| $AIC = -136$ ; $RMSE = .05$ ; $NRMSE = .80$ ; $F(4,40) = 4.6$ , $p = .004^{**}$ ; $\Delta R^2 = .25$ |  |  |  |  |  |  |  |
| CB mod (45; 8.8) | **THAL** | **6.3** | **1.8** | **3.5** | **.55** | **2.75, 9.85** | **< .01 \*\*** |
|  | PUT | -3.6 | 2.1 | -1.71 | -.22 | -7.7, -.52 | = .09 |
|  | HPC | 1.8 | 1.55 | 1.15 | .17 | -1.2, 4.8 | = .25 |
| $AIC = 106$ ; $RMSE = .70$ ; $NRMSE = .76$ ; $F(3,42) = 9.2$ , $p < .0001^{***}$ ; $\Delta R^2 = .36$ |  |  |  |  |  |  |  |
| LV mod<sup>T</sup> (45; 2.4) | **THAL** | **-3.08** | **.461** | **-6.68** | **-.69** | **-3.9, -2.2** | **< .0001 \*\*\*** |
|  | **Age** | **1.47** | **.007** | **3.1** | **.28** | **.45, 2.5** | **< .01 \*\*** |
| $AIC = 35$ ; $RMSE = .33$ ; $NRMSE = .61$ ; $F(2,42) = 34.1$ , $p < .0001^{***}$ ; $\Delta R^2 = .61$ |  |  |  |  |  |  |  |
| THAL mod 1 (45; .72) | **PUT** | **.313** | **.132** | **2.4** | **.22** | **.05, .57** | **= .023\*** |
|  | **HPC** | **.213** | **.097** | **2.2** | **.23** | **.02, .40** | **= .034\*** |
|  | **LAT VENT** | **-.0235** | **.006** | **-3.9** | **-.44** | **-.04, -.01** | **= .0003 \*\*\*** |
|  | **CB** | **.023** | **.0095** | **2.4** | **.26** | **.00, .04** | **= .02\*** |
| $AIC = -139$ ; $RMSE = .05$ ; $NRMSE = .56$ ; $F(4,40) = 21.2$ , $p < .0001^{***}$ ; $\Delta R^2 = .65$ |  |  |  |  |  |  |  |
| THAL mod 2 (45; .71) | **PUT** | **.29** | **.13** | **2.2** | **.21** | **.04, .56** | **= .031\*** |
|  | **HPC** | **.22** | **.094** | **2.3** | **.24** | **.03, .40** | **= .025\*** |
|  | **HPC: Age** | **-.031** | **.0144** | **-2.14** | **-.25** | **-.06, -.01** | **= .04\*** |
|  | **CB** | **.026** | **.009** | **2.8** | **.30** | **.008, .04** | **= .009 \*\*** |
|  | **LAT VENT** | **-.02** | **.006** | **-3.2** | **-.38** | **-.03, -.01** | **= .002 \*\*** |
|  | Age | -.0015 | .0012 | -1.2 | -.13 | -.004, 0 | = .23 |

$AIC = -140$ ; $RMSE = .04$ ; $NRMSE = .53$ ; $F(6,38) = 13.1$ , $p < .0001^{***}$ ; $\Delta R^2 = .67$
$\Delta R^2$ = adjusted $R^2$ $AIC$ = Akaike's Information Criterion; $\beta$ = standardized coefficient; Effect = variable name; CB:Age = cerebellar by age interaction higher order term; Est = coefficient; F = F-statistic; HPC = hippocampal normalized volume; LV = lateral ventricle; mod = model; NRMSE = normalized RMSE; PUT = putamen normalized volume; THAL = normalized thalamic volume; CB = normalized cerebellar volume; <sup>T</sup> = log transformed dependent and explanatory variables. All values are rounded. All continuous predictors were centered in the THAL mod 2, which included a HPC by age interaction; all continuous predictors were also centered in the PUT model, which included an CB by age interaction.

A non-linear relation found initially in the lateral ventricular model was corrected by log transform of the dependent and independent variables. All final models complied with assumptions as tested using a comprehensive test (^63^) and plots (see *Supporting information I*, S1-2 to S1-12). There were no overly influential cases as measured by Cook’s distance (4/n). Two thalamic volume models were retained because one of them (THAL mod 2, see **Table 3**), included an interaction term. This precluded compliance with the ten (data instances) to one predictor rule of thumb (^58^), a guideline to mitigate against overfitting. The THAL mod 2, while not complying with this rule of thumb, provides some insight into a hippocampal volume by age interaction that could occur in a larger data set. Moreover, the THAL mod 2 did comply with the less a stringent one predictor to five data instance rule of thumb (^57^). In addition, as a further check against overfitting, a predictive R-squared calculation was applied to THAL mod 2. There was only a small discrepancy between the THAL mod 2 adjusted *R^2^*.67 and the predictive *R^2^* outcome of .62, suggesting that this model was not overfitted. Nevertheless, a thalamic model compliant with the traditional ten to one rule of thumb (THAL mod 1, **Table 3**) was also retained.

In the hippocampal volume model HPC mod 1, stepwise regression (backward, using AIC), used to help narrow the final set of all model predictors, originally retained the explanatory variable cerebellar (CB) volume. The CB predictor was not retained as it made negligible difference to the hippocampal (HPC mod 1) model fit (based on the RMSE and *adjusted-R**^2^***). In the CB (cerebellar volume as the response variable) and putamen (putamen volume as the response variable) volume models, insignificant predictors were retained because they contributed more to model fit of the data (based on the RMSE and *adjusted-R**^2^***).

FIG 3 effects plots are for each model’s most important continuous predictors. Effects plots use partial residuals, meaning that while each plot is derived from a complete model (i.e., the models in **Tables 3**), variables other than the effect of interest plotted are controlled for (^64^).

**FIG 3:**
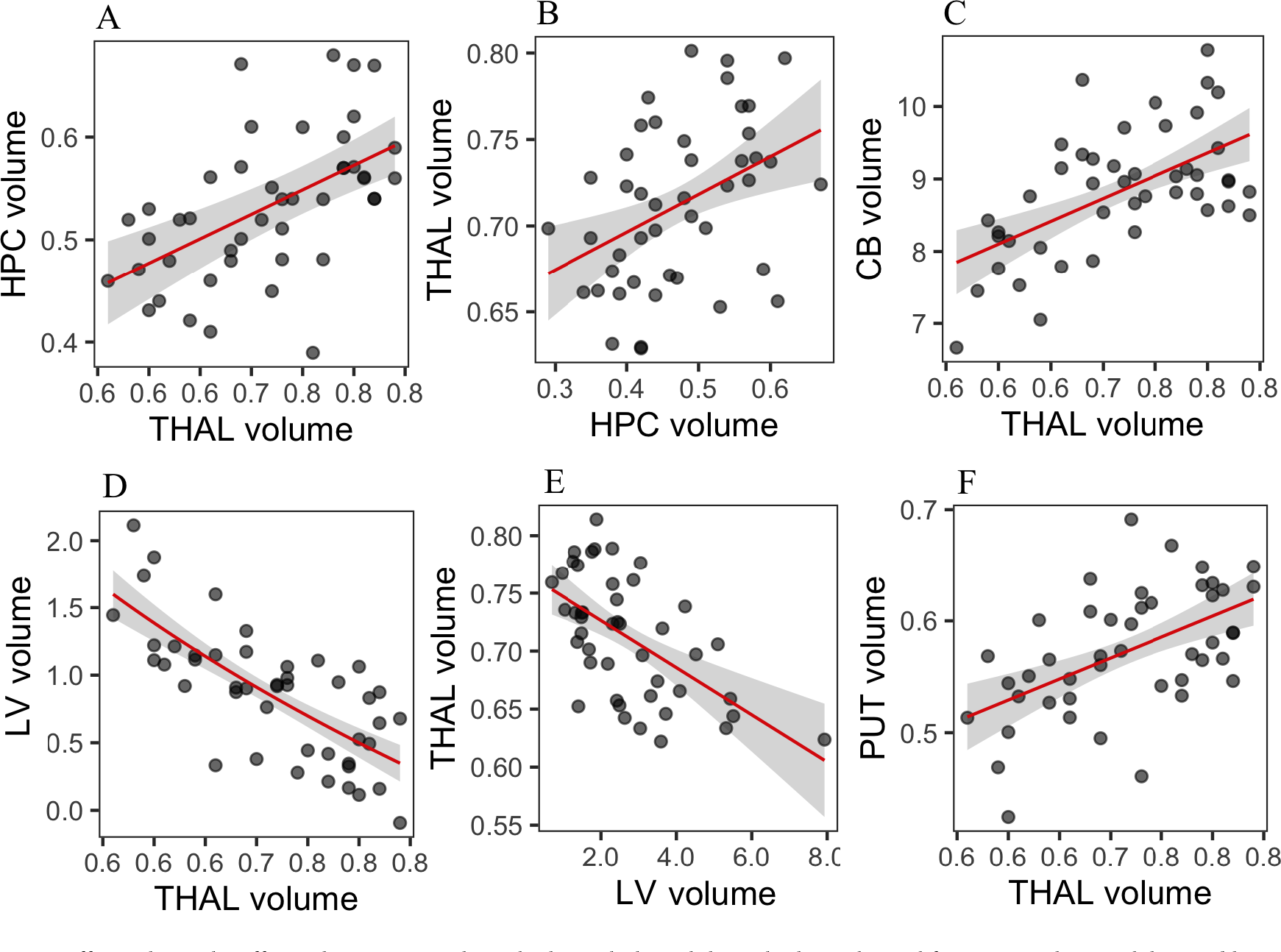
Effects plots: The effects plots use partial residuals, such that while each plot is derived from a complete model, variables other than the effect of interest plotted are controlled for. A = hippocampal (HPC) on thalamic volume, B = thalamic (THAL) on HPC volume, C = cerebellar (CB) on THAL volume, D = lateral ventricular (LV) on THAL volume, E = thalamic (THAL) on LV volume, F *= putamen (PUT) on THAL volume. All volume measures are normalized and relative as a % total intracranial volume*.

FIG 3 includes two effects plots for the thalamic volume model: thalamic volume on hippocampal volume (FIG 3B), and thalamic volume on lateral ventricular volume (FIG 3E). Thalamic volume as an explanatory variable had consistently the greatest effect. Four of six plots in FIG 3 help to convey the explanatory effect of thalamic volume in linear models, where volume of one of the other structures was the response variable.

### 4.2 Model interpretation

In brief example interpretations, with respect to the HPC mod 1, **Table 3**, controlling for group (AD and MCI group membership), a unit increase in thalamic volume explained a .48% increase (.48% of intracranial volume [ICV]) in hippocampal volume that was statistically significant, *p* = .0003. Holding thalamic volume as well as the MCI group membership constant, AD group membership explained a .1% decrease in mean hippocampal volume relative to controls, a difference that significantly differed from zero, *p* < .0001. Holding thalamic volume as well as AD group membership constant, MCI group membership explained a .08% decrease in mean hippocampal volume relative to controls, a difference that significantly differed from zero, *p* = .001. Inputting the grand mean thalamic volume of .72% into the model (HPC mod 1), the model estimated hippocampal volume marginal means (means predicted by the fitted model) were as follows: AD .43% (95% CI .40, .46), MCI .45% (95% CI .42, .48), CN.52% (95% CI .50, .56). A Tukey test indicated mean hippocampal volume was significantly reduced in AD (*M*=. 43%, *SD*=. 08) versus controls (*M*=. 52%, *SD*=. 07), *p* <. 001, *d*= -1.6, and in MCI (*M*=. 45% *SD*=.06) versus controls (*M* =. 52%, *SD*=. 07), *p* <.01, *d* = -1.6. There was no significant difference between AD and MCI model means, *p* > .05. Note also that a binarized version (see Methods) of cognitive dementia rating (CDR) had a similar effect to the group variable (AD vs. control) explaining hippocampal volume.

With regard to log transformed(^52^) lateral ventricular model, holding thalamic volume constant, for every 10% increase in age (range 56 to 88) lateral ventricular volume increased by about 15% (of total intracranial volume [ICV]), which significantly differed from zero, t(42) = 2.8, *p* =.007; holding age constant, every 10% increase in thalamic volume explained a lateral ventricular volume decrease of about 25% (of ICV), t(42) = -6.68, *p* < .0001. The latter reflects, for the most part, the anatomical thalamus-lateral ventricular juxtaposition: the superior aspect of the thalamus forms much of lateral ventricular inferior border or floor (^65^). The comprehensive test suite (^63^) for assumptions indicated no violations of assumptions. Converting the model log coefficients back to original values, inputting the average age of 73.3 and grand mean for thalamic volume of .72%, the predicted lateral ventricular mean volume (based on uncentered predictors in this case) was 2.4% (of ICV) (95% CI 2.1, 2.6).

### 5.1 Bivariate repeated-measures analysis

**Table 4** summarizes the repeated measures strength and direction of association between structure volume and a given predictor. All values represent bivariate repeated-measures correlations (*rr*_m_) ignoring groups. As noted in section 2.5, the *rr*_m_ coefficients have a -1 to 1 index of between variable association strength analogous to Pearson *r*, but unlike Pearson *r* the *r*_rm_ calculation does account for the non-independence of repeated-measures (^43^).

**TABLE 4:** Bivariate repeated-measure correlations (rrm), N=36 (108 data points over 3 time points)

| Variable | HPC Vol | PUT Vol | THAL Vol | LAT VENT Vol | CB Vol |
| --- | --- | --- | --- | --- | --- |
| | $r_{rm}$ (df),<br>95% CI (),<br>p-value | $r_{rm}$ (df),<br>95% CI,<br>p-value | $r_{rm}$ (df),<br>95% CI,<br>p-value | $r_{rm}$ (df),<br>95% CI,<br>p-value | $r_{rm}$ (df),<br>95% CI,<br>p-value |
| **HPC Vol** | | $r_{rm}(71) = .45$<br>(.24, .62),<br>$p < .0001$ | $r_{rm}(71) = .74$<br>(.61, .83),<br>$p < .0001$ | $r_{rm}(71) = -.68$<br>(-.79, -.54),<br>$p < .0001$ | $r_{rm}(71) = .22$<br>(-.01, .43),<br>$p = .06$ |
| **PUT Vol** | | | $r_{rm}(71) = .60$<br>(.42, .73),<br>$p < .0001$ | $r_{rm}(71) = -.39$<br>(-.57, -.17),<br>$p < .001$ | $r_{rm}(71) = .26$<br>(.03, .46),<br>$p = .03$ |
| **THAL Vol** | | | | $r_{rm}(71) = -.55$<br>(-.68, -.36),<br>$p < .0001$ | $r_{rm}(71) = .001$<br>(-.23, .23),<br>$p = .99$ |
| **LAT VENT Vol** | | | | | $r_{rm}(71) = -.43$<br>(-.60, -.22),<br>$p < .001$ |
| **Age** | $r_{rm}(71) = -.63$,<br>(-.76, -.47),<br>$p < .0001$ | $r_{rm}(71) = -.27$,<br>(-.47, -.04),<br>$p = .02$ | $r_{rm}(71) = -.46$,<br>(-.62, -.25),<br>$p < .0001$ | $r_{rm}(71) = .75$,<br>(.63, .84),<br>$p < .0001$ | $r_{rm}(71) = -.30$,<br>(-.50, -.07),<br>$p = .009$ |
| **CDR** | $r_{rm}(71) = -.43$,<br>(-.6, -.22),<br>$p < .001$ | $r_{rm}(71) = -.23$<br>(-.44, .001),<br>$p = .048$ | $r_{rm}(71) = -.33$,<br>(-.52, -.1),<br>$p = .005$ | $r_{rm}(71) = .46$,<br>(.26, .63),<br>$p < .0001$ | $r_{rm}(71) = -.13$,<br>(-.35, .11),<br>$p = .27$ |
| MMSE | $r_{rm}(71) = .32, (.09, .51),$<br>$p < .01$ | $r_{rm}(71) = 0.22,$<br>(0.04 0.47),<br>$p = .055$ | $r_{rm}(71) = 0.27,$<br>(.03, .47),<br>$p = .02$ | $r_{rm}(71) = -.36,$<br>(-.54, -.14),<br>$P = .002$ | $r_{rm}(71) = 0.25,$<br>(.02, .46),<br>$p = .03$ |

CB = cerebellum; HPC= hippocampus; PUT = putamen; THAL = thalamus; LAT VENT = lateral ventricles; $r_{rm}$ = repeated-measures correlation coefficient (Bakdash, Marusich, 2017); Vol = normalized volume. Strong effect correlations ( $r_{rm} \geq .60$ ) are in bold

### 6.1 Longitudinal analysis

This section reviews the mixed model analyses for longitudinal volume analysis of hippocampal, putamen, thalamic, cerebellar, and lateral ventricular structures across the three time points: baseline, 12 months and 24 months. See section *2.5.2* *Linear mixed model methods* for methodological details. Longitudinal findings begin with a short summary of annualized volume results. This is followed by graphs, then the tabulated mixed model details are reviewed. Finally, an example model interpretation is provided focusing on the longitudinal hippocampal volume model. Additional model details, including comparisons of each model type (1-7: see *2.5.2 Linear mixed model methods*) are provided in *Supporting information 1*. Note, in aid of reducing this article length, mixed model diagnostics, which included fixed and random effects assessments, are provided in *Supporting information VI*. All final models complied with mixed model regression assumptions.

Note, some subjects did not have measures at 12- and 24-month times. In addition, to comply with model diagnostic findings (see *Supporting information VI*) a few more data instances were removed (see **Tables 5, 6** for each structure’s model *n*-count). To reduce risk of overfitting, all mixed models complied with the relaxed one predictor to five data instance rule of thumb (^57^).

**TABLE 5:**
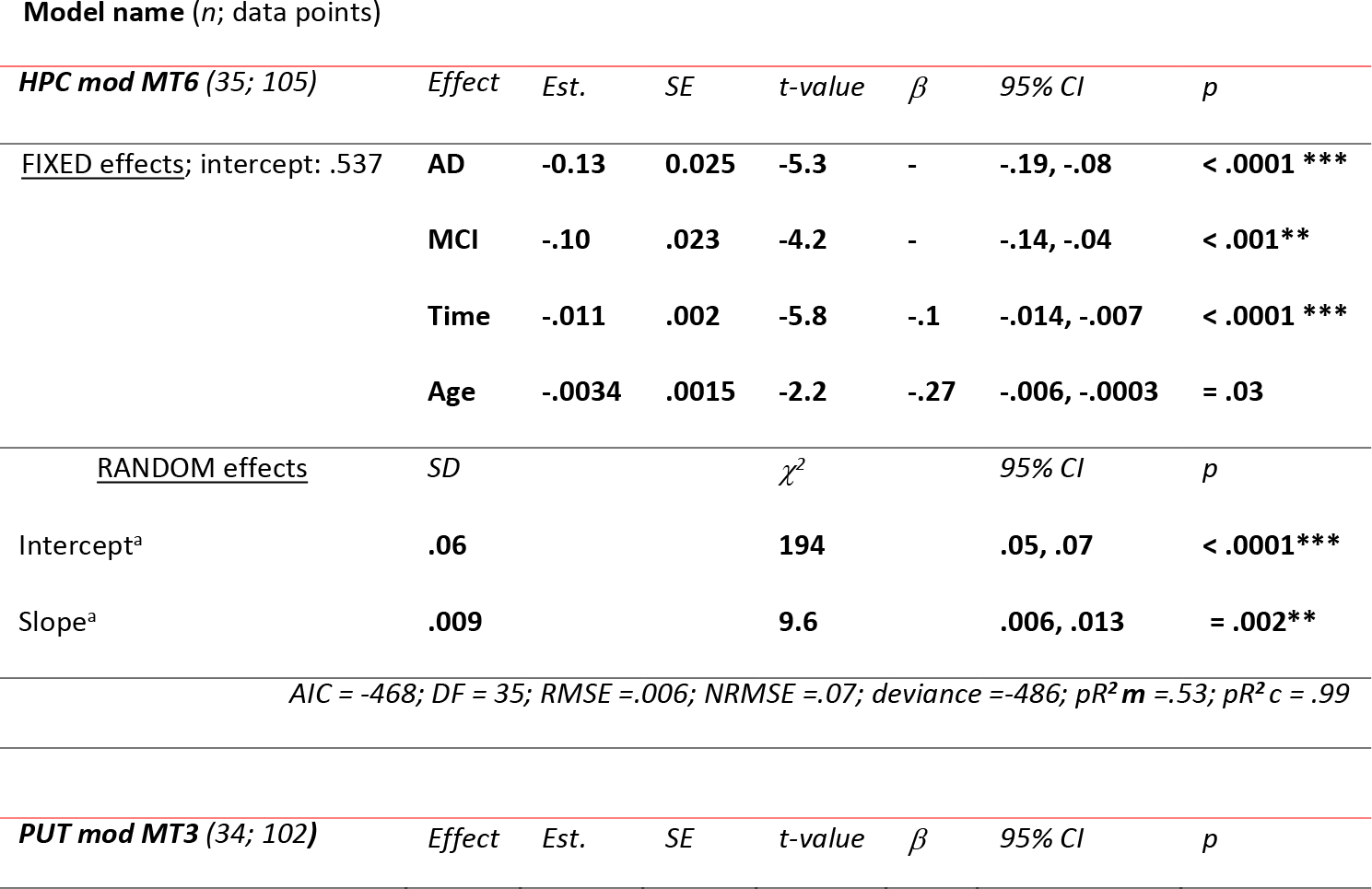
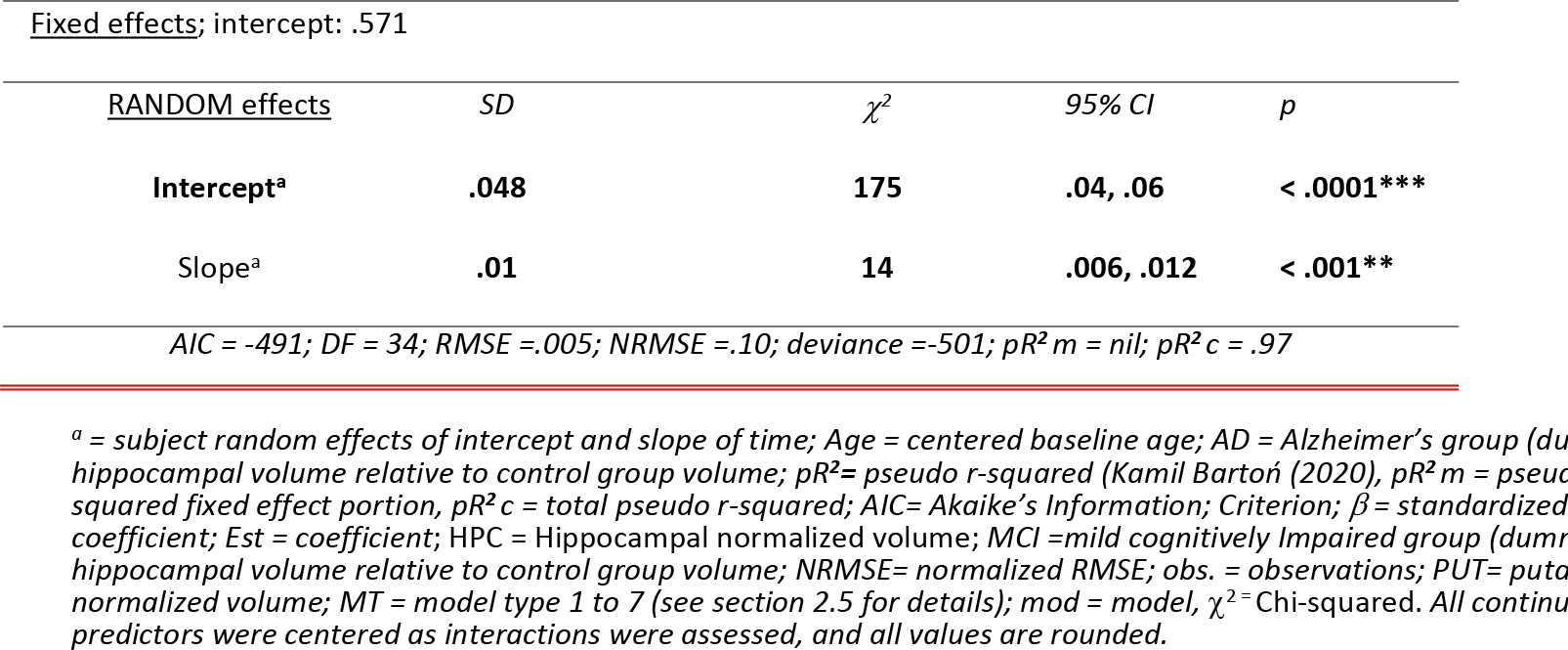
Mixed models: hippocampal and putamen volume

**TABLE 6:**
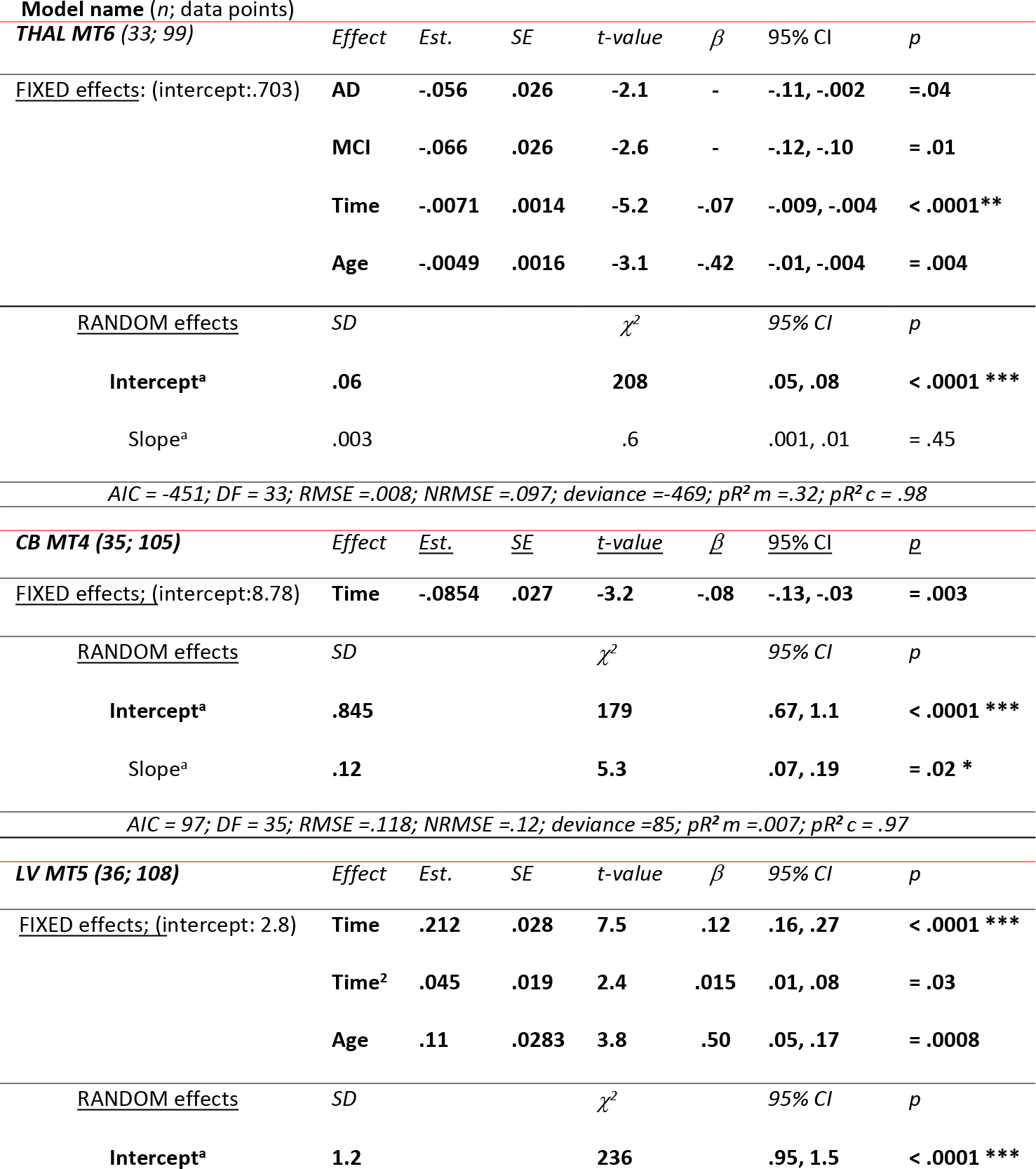
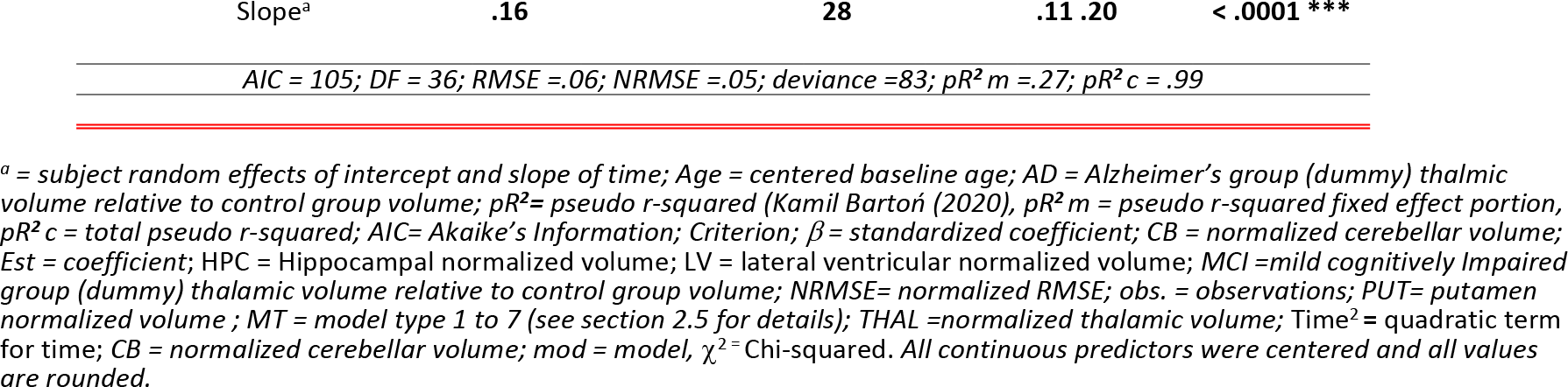
Mixed models: thalamic, cerebellar and lateral ventricular volume

| Model name (*n*; data points) |  |  |  |  |  |  |  |
| --- | --- | --- | --- | --- | --- | --- | --- |
| **THAL MT6 (33; 99)** | *Effect* | *Est.* | *SE* | *t-value* | $\beta$ | *95% CI* | *p* |
| <u>FIXED effects</u>: (intercept: .703) |  |  |  |  |  |  |  |
|  | AD | -.056 | .026 | -2.1 | - | -.11, -.002 | = .04 |
|  | MCI | -.066 | .026 | -2.6 | - | -.12, -.10 | = .01 |
|  | Time | -.0071 | .0014 | -5.2 | -.07 | -.009, -.004 | < .0001** |
|  | Age | -.0049 | .0016 | -3.1 | -.42 | -.01, -.004 | = .004 |
| <u>RANDOM effects</u> | *SD* | | | $\chi^2$ | | *95% CI* | *p* |
| Intercept<sup>a</sup> | .06 |  |  | 208 |  | .05, .08 | < .0001 *** |
| Slope<sup>a</sup> | .003 |  |  | .6 |  | .001, .01 | = .45 |
| AIC = -451; DF = 33; RMSE = .008; NRMSE = .097; deviance = -469; $pR^2 m = .32$; $pR^2 c = .98$ | | | | | | | |
| **CB MT4 (35; 105)** | *Effect* | *Est.* | *SE* | *t-value* | $\beta$ | *95% CI* | *p* |
| <u>FIXED effects</u>: (intercept: 8.78) |  |  |  |  |  |  |  |
|  | Time | -.0854 | .027 | -3.2 | -.08 | -.13, -.03 | = .003 |
| <u>RANDOM effects</u> | *SD* | | | $\chi^2$ | | *95% CI* | *p* |
| Intercept<sup>a</sup> | .845 |  |  | 179 |  | .67, 1.1 | < .0001 *** |
| Slope<sup>a</sup> | .12 |  |  | 5.3 |  | .07, .19 | = .02 * |
| AIC = 97; DF = 35; RMSE = .118; NRMSE = .12; deviance = 85; $pR^2 m = .007$; $pR^2 c = .97$ | | | | | | | |
| **LV MT5 (36; 108)** | *Effect* | *Est.* | *SE* | *t-value* | $\beta$ | *95% CI* | *p* |
| <u>FIXED effects</u>: (intercept: 2.8) |  |  |  |  |  |  |  |
|  | Time | .212 | .028 | 7.5 | .12 | .16, .27 | < .0001 *** |
|  | Time<sup>2</sup> | .045 | .019 | 2.4 | .015 | .01, .08 | = .03 |
|  | Age | .11 | .0283 | 3.8 | .50 | .05, .17 | = .0008 |
| <u>RANDOM effects</u> | *SD* | | | $\chi^2$ | | *95% CI* | *p* |
| Intercept<sup>a</sup> | 1.2 |  |  | 236 |  | .95, 1.5 | < .0001 *** |

|  |  |  |  |  |
| --- | --- | --- | --- | --- |
| Slope<sup>a</sup> | .16 | 28 | .11 .20 | < .0001 *** |
| AIC = 105; DF = 36; RMSE = .06; NRMSE = .05; deviance = 83; pR<sup>2</sup> m = .27; pR<sup>2</sup> c = .99 |  |  |  |  |

<sup>a</sup> = subject random effects of intercept and slope of time; Age = centered baseline age; AD = Alzheimer's group (dummy) thalamic volume relative to control group volume; pR<sup>2</sup> = pseudo r-squared (Kamil Bartoń (2020), pR<sup>2</sup> m = pseudo r-squared fixed effect portion, pR<sup>2</sup> c = total pseudo r-squared; AIC = Akaike's Information Criterion; β = standardized coefficient; CB = normalized cerebellar volume; Est = coefficient; HPC = Hippocampal normalized volume; LV = lateral ventricular normalized volume; MCI = mild cognitively impaired group (dummy) thalamic volume relative to control group volume; NRMSE = normalized RMSE; obs. = observations; PUT = putamen normalized volume; MT = model type 1 to 7 (see section 2.5 for details); THAL = normalized thalamic volume; Time<sup>2</sup> = quadratic term for time; CB = normalized cerebellar volume; mod = model, χ<sup>2</sup> = Chi-squared. All continuous predictors were centered and all values are rounded.

The assessment of median annualized percentage volume change over a period of two years was completed subsequent to the mixed model analysis, and provides another perspective of volume change over time. To be consistent with the mixed model findings, this calculation (see *Statistics 2.5*) included a group breakdown only for hippocampal and thalamic annualized percentage volume change calculations. The best fitting hippocampal and thalamic mixed models, as reviewed below, retained the group (AD, CN, MCI) variable; the best fitting cerebellar, putamen and lateral ventricular mixed models, however, did not retain the group variable, and annualized percentage volume change for these structures was assessed without specifying group. The hippocampal and thalamic median annualized percentage volume change (by group) clearly followed the AD > MCI > CN pattern of greater annualized volume reduction. Hippocampal median annualized volume change (by group) ranged from -4.7% in AD to -.88% in controls. Median annualized thalamic volume change (by group) ranged from -1.3% in AD to -.62% in controls. The heightened hippocampal volume change in AD amounted to about 2.5 times the annual reduction in MCI; change in MCI, reduction in hippocampal volume, amounted to 2 times that found in controls. The extent of thalamic volume reduction in AD was about 1.6 times the annual reduction in MCI and about 2 times the annual reduction that occurred in controls. The median annualized percentage change, not specified by group, for cerebellar, putamen and lateral ventricular volume was -.60, 0, and 6.8 respectively. Details are provided in *Supporting Information II* (see Tables S2-1 and S2-2).

FIGs 4 and 5 reveal the longitudinal (2-year) pattern in the volumetric data. FIG 4 (Trellis) plots, panels A to E, are based on 108 data instances, 36 at each time point (see Table 2 for the group breakdown). They convey the trend in structure volume change over time for each subject. Specifically, the random, individual subjects’ intercepts (volume means) from baseline to 24 months are provided; individual structure volume is regressed on time. The upper panel area in each of these plots has the subject anonymized identification number; a regression line in each subject’s plot shows the slope (ordinary least squares estimated line for a given subject panel), which, here, is the subject volume extent of change at each time-point, and the intercept representing the average, normalized volume for each subject. Close inspection of FIG 4 shows hippocampus (A) and thalamic (B) volume with largely linear decreases in volume over time.

**FIG 4.**
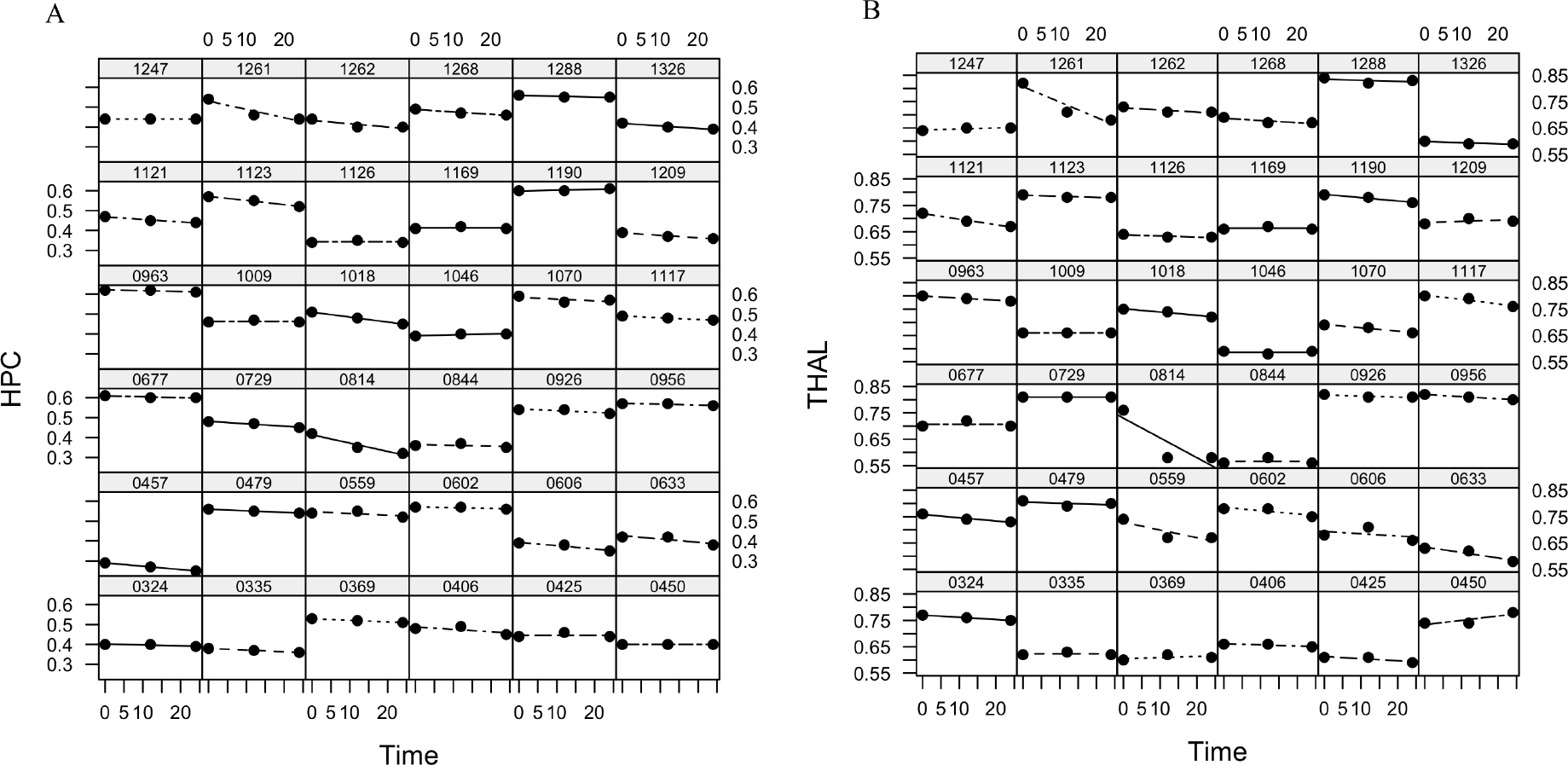
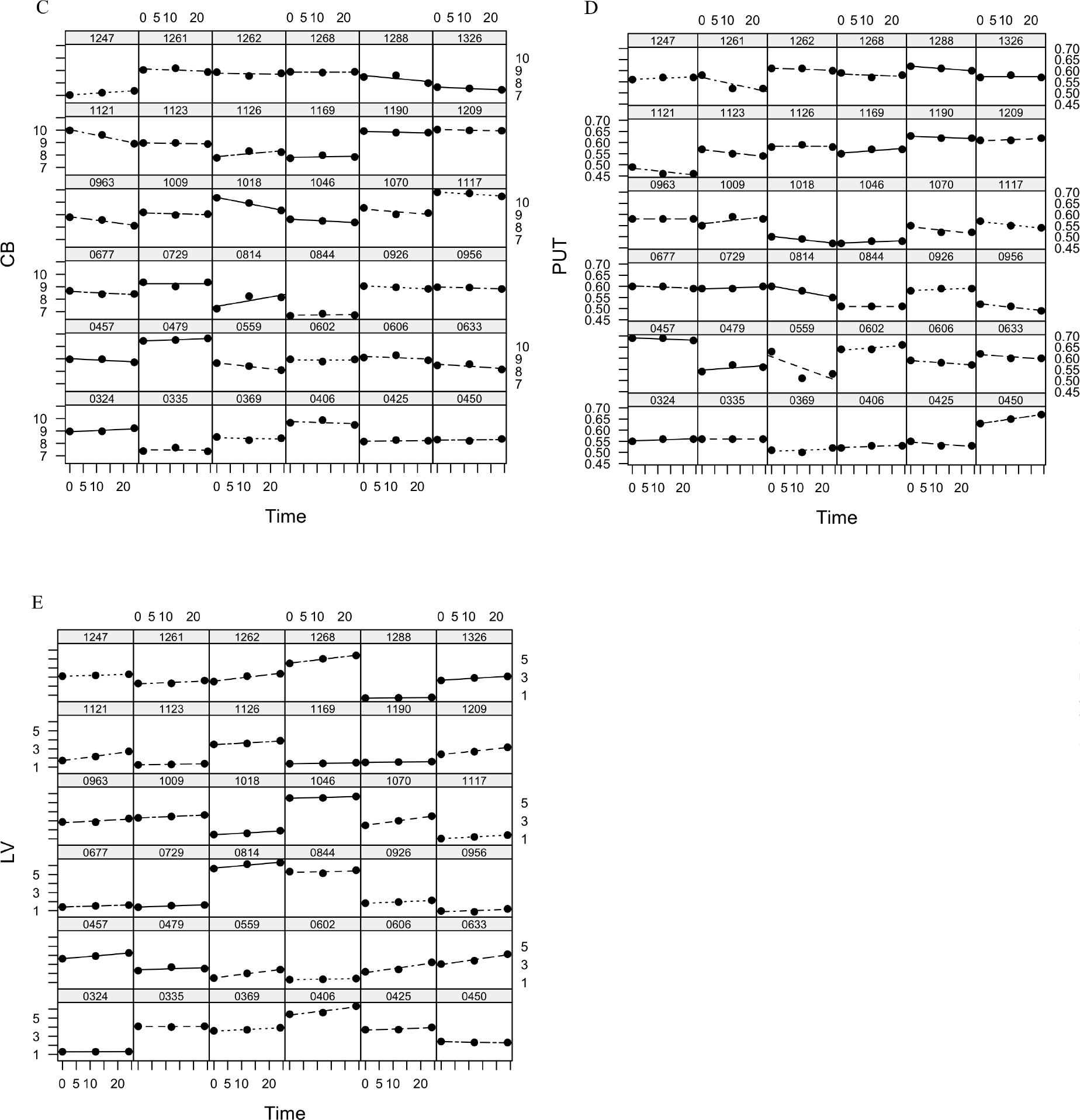
Lattice plots: Individual subject intercepts (volume) and time trends. A = hippocampus (HPC); B = thalamus (THAL); C = cerebellum (CB); D = putamen (PUT); E =lateral ventricles (LV). Y-axis is the structure’s percentage of total intracranial volume, plotted against time: 0 to 24 months. Numbers inside and at the top of each panel are subject identifiers. Data for each subject are provided in an individual panel along with a simple linear regression line. Considerable variation of random subject intercepts over time is evident.

The cerebellum (C) has a similar pattern of volume decrease over time and the putamen (D) has a less distinct pattern of volume change over time. The lateral ventricle (E) volume generally increases linearly over time.

Longitudinal baseline (0) to 24-month (spaghetti) plots are provided in FIG 5. Thicker lines are the group mean. The thick group mean lines help to clarify patterns: e.g., note the largely antithetical hippocampal (A) and lateral ventricular (E) volume change patterns particularly evident in the AD group.

**FIG 5:**
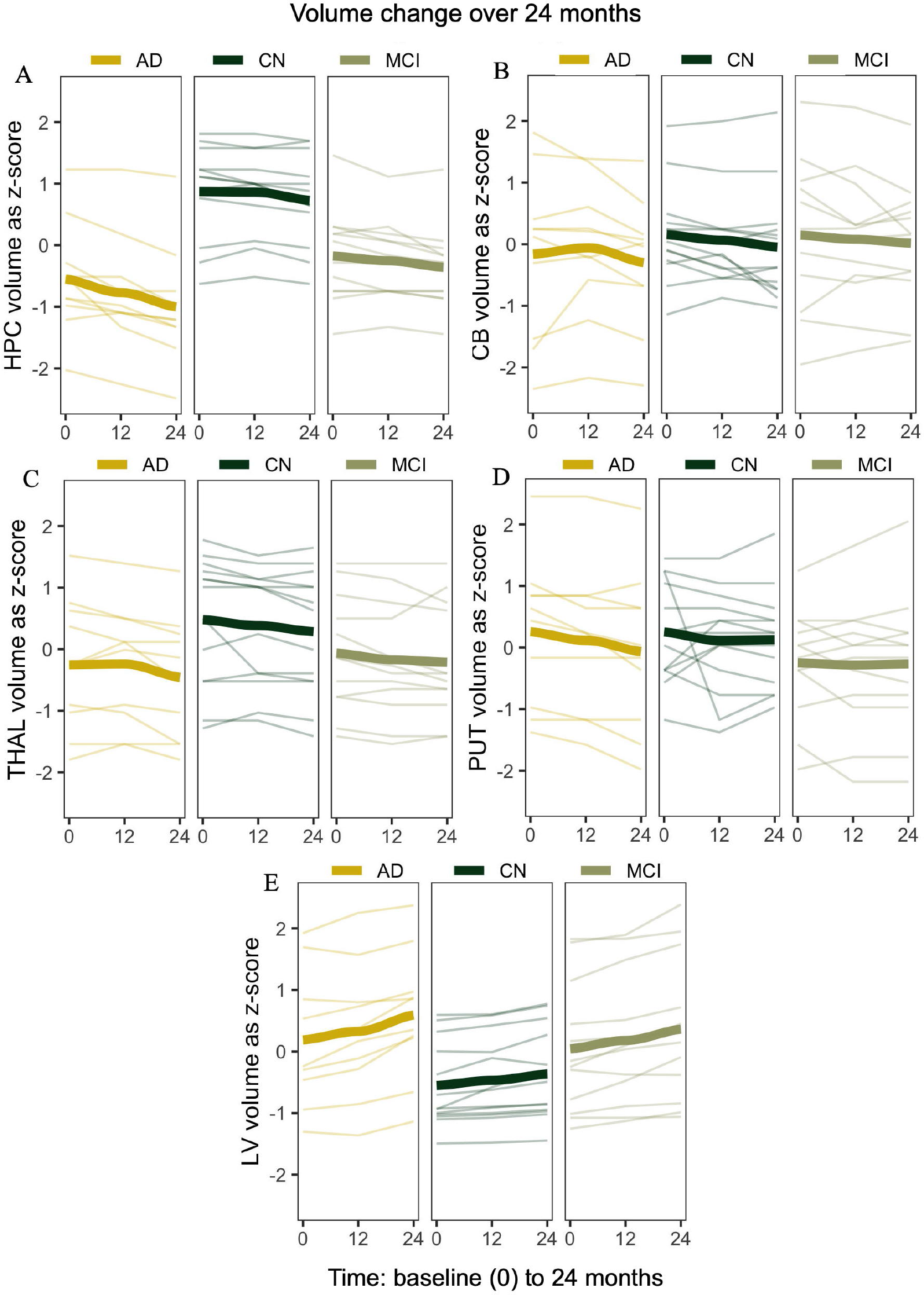
Spaghetti plots: Y-axes show normalized volume converted to z-scores; x-axes show time, baseline to 24 months (baseline = 0). (A) hippocampus (HPC); (B) cerebellum (CB); (C) thalamus (THAL); (D) putamen (PUT); (E) lateral ventricles (LV). AD = Alzheimer’s, CN = controls, MCI = mild cognitively impaired. Thicker lines are group means.

### 6.2 Mixed model results

The mixed longitudinal models examined change, if any, in structure volume related to the fixed effects of passage of time (baseline, 12 months, 24 months), baseline age, and group (AD, CN, MCI) membership. Random effects assessed were the subject intercept (equivalent to individual subject variation of intercept) and the subject random slope of time (an indication of extent the effect of time varies per subject). Details are tabulated in **Tables 5** and **6**. The normalized volume of each structure broken down by group and time was provided in **Table 2**. A quadratic term for the fixed effect of time was initially included in models of each structure. However, a quadratic term significantly contributed to only the lateral ventricular model and was only retained in the lateral ventricular model. A version of the hippocampal volume mixed model included an interaction (see *Supporting Information I*, Table S1-1). However, this interaction was not tabulated in the results section because there were too few data instances to formally support this more complex model.

An example interpretation is provided in section 6.3, which focuses on the hippocampal mixed model results. The hippocampal model predicted means are also provided in section 6.3. The predicted thalamic volume means at baseline mean age and at all three time-points were as follows for each group. Controls: baseline time, .746 (95% CI .705, .786); 12 months,.739 (95% CI .699, .779); 24 months, .732 (95% CI .692, .771). MCI: baseline time, .679 (95% CI .637, .722); 12 months,.672 (95% CI .630, .714); 24 months, .665 (95% CI .623, .707). AD: baseline time, .690 (95% CI .645, .734); 12 months, .682 (95% CI .638, .727); 24 months, .675 (95% CI .632, .719). Here, all confidence intervals overlap, and while this is often an indication of non-significant group differences, such is not always the case. Indeed, pairwise tests indicated that the MCI and control group marginal means significantly differed as determined by a Tukey test, z = -2.6, *p* = .03, *d* = -.83. AD vs. MCI did not significantly differ (z = -0.38*, p* =.92, *d* = -.63), nor did AD vs. controls (z = -2.12, *p* = .09, *d* = -.74).

The model predicted volume means for the cerebellum, putamen, and lateral ventricles were as follows. The cerebellum, in addition to random effects, retained only the fixed effect of time. The predicted cerebellar volume mean was 8.87% (95% CI 8.6, 9.2) at baseline time, 8.79% (95% CI 8.5, 9.1) at 12 months, and 8.70 (95% CI 8.4, 9) at 24 months. The putamen model predicted mean (amounting to the intercept in this case given no fixed effects were retained in the best model, just random effects) was .57 (95% CI .55, .59). The lateral ventricular best fitting model retained the fixed effect of time, coupled with a time quadratic term (time^2^), and baseline age (in addition to the random effects, but not group). Model predicted means were 2.60% (95% CI 2.2, 3.0) at baseline, 2.84% (95% CI 2.43, 3.26) at 12 months, and 3.07% (95% CI 2.63, 3.51) at 24 months.

### 6.3 Hippocampal mixed model analysis and interpretation

The final saturated model, model type 6 in this case, included fixed effects of time, baseline age, group (AD, CN, MCI), and random effects of subject intercept and time. Subject random effects remained significant. This indicates that individual differences (random effects) were not completely accounted for by the fixed effects (time and group). As indicated in the random effects portion of the hippocampal model (HPC mod MT6) **Table 5**, over time, both the subject means (the intercept) and slope of time (varying effect of time) remained significant in the saturated model type 6. Comparisons of the model types 1-7 (based on AIC, BIC, loglikelihood and p-values) is provided in Supporting information I (Table S1-1).

Hippocampal mixed model results in **Table 5** is consistent with FIGs 2 (A) and 5 (A). A greater decrease in volume over time is evident in FIG 5 for AD and MCI groups relative to controls. Also, conveyed in FIG 5 (A), hippocampal (normalized) volume against time was not constant but appeared to depend on group, indicating that the volume change per each 12-month interval (across baseline, 12 months, and 24 months) varied by group (an interaction). A significant (*p* = .003) AD group by time higher order term is reported in Table S1-2 (model 7 type, see *Supporting information I*). However, as already noted, the sample size was too small to reliably support the inclusion of an additional term, here the higher order term of the model 7 type derived from the group by time interaction. As such, a less complex additive model 6 type is reported in **Table 5**. Notwithstanding the potential of an interaction persisting in a larger data set, FIG 2 (A) and FIG 5 (A) indicate the control and MCI hippocampal volume means were, at each time-point for most subjects, above the AD mean. This is one indication that outcome was not driven by an interaction, and the main effects (HPC mod MT6, **Table 5**) warrant comment.

Elaborating briefly on the main effects of model type 6 (no interaction), controlling for group, the overall effect of a year’s passage of time, on average, significantly predicted a 0.011% (95% CI -.014, -.007) decrease (:: .01% of ICV) per year in hippocampal volume, *t*(35) = -5.8, *p* < .0001. Controlling for time, baseline age and MCI group membership, AD group membership, relative to controls, significantly predicted a 0.13% (95% CI -.19, -.08) decrease in hippocampal volume, *t*(35) = -5.3, *p* < .0001. Controlling for time, baseline age, and AD group membership, MCI group membership, relative to controls, significantly predicted a .10% volume (95% CI - 0.14, -0.04) decrease, *t*(35) = -4.2, *p* < .001. Finally, controlling for other predictors, baseline age significantly predicted a .003% (95% CI -.006, -.0003) decrease in hippocampal volume, *t*(35) = -2.2, *p* = .03. Lending some perspective to the standardized coefficients, and using baseline age as an example, there was a .27 standard deviation reduction in hippocampal volume per year of age. This constitutes about a .02% (*SD* hippocampal volume = .087; .27 * .087 = .02) decrease per year of hippocampal volume as derived from baseline age. The marginal (fixed effect *pR^2^m*) pseudo *R^2^*^(^<u>^54^</u>^)^ for this model was .53 (i.e., 53% of pseudo variance, not actual variance, was explained by the model) and the composite fixed and random effects (*pR^2^c*) was .99. The RMSE and NRMSE for this longitudinal hippocampal volume analysis were relatively small and the pseudo *R^2^*for the fixed effects (*pR^2^m)* was relatively high.

The group predicted marginal means (average fitted marginal, by group, means) of hippocampal volume at mean baseline age and at all three time-points were as follows for each group. Controls: baseline time, .548 (95% CI .511, .584); 12 months,.537 (95% CI .501, .573); 24 months, .526 (95% CI .490, .563). MCI: baseline time, .446 (95% CI .408, .484); 12 months,.436 (95% CI .398, .474); 24 months, .425 (95% CI .387, .463). AD: baseline time, .413 (95% CI .411, .455); 12 months,.402 (95% CI.361, .444); 24 months, .392 (95% CI .350, .433).

Note, the confidence intervals for AD and controls do not overlap, nor the confidence intervals for controls and MCI; one indication that both AD and MCI groups significantly differ from controls. On the other hand, the AD and MCI confidence intervals do overlap, an indicator of lack of significant difference between AD and MCI hippocampal volumes.

In post hoc pairwise comparisons using a Tukey test, AD members had significantly lower hippocampal volume compared to controls, z = -5.3, *p* < .0001, *d* = -1.9. This *d*-value indicates a large negative effect of AD group membership on hippocampal volume. MCI hippocampal volume also significantly differed from controls, z = -4.2, *p* < .0001, *d* = -1.56. Again, as with the AD group, this d-value indicated MCI group membership had a large negative effect on hippocampal volume. There was no significant hippocampal volume difference between AD and MCI groups, z = 1.2, *p* =.23 *d* = -.62.

## Discussion

Utilizing subcortical structure volume as the primary measure of interest in AD and normal aging, the current work was a composite study uniquely combining three phases of research. Initially, a platform dedicated to volumetry analysis, volBrain(^42^), was used for the automated segmentation/volume calculation of five structures: cerebellum, putamen, hippocampus, lateral ventricles and thalamus. After a reliability assessment of volBrain volume estimates, linear regression and linear mixed longitudinal regression analyses were conducted.

The test-retest reliability of volBrain segmented same-subject scans, was high, as indicated by the mean ICC score of .989 (*SD* = .012). The lowest ICC was .937 (95% CI .829, .978). For details on all ICC measures see **Table** S1-11 in *Supporting Information I*.

There were a few salient, notable findings. First, linear models demonstrated thalamic volume had consistently the greatest explanatory effect of volume in the other structures: the hippocampus, putamen, cerebellum and lateral ventricles. Second, in longitudinal mixed models, the group variable AD and MCI cohorts (that served as proxies for early AD and MCI respectively) significantly contributed to the best-fitting hippocampal and thalamic models but not to the best fitting putamen, cerebellar or lateral ventricular models. Third, longitudinal mixed models indicated time (1 year) had a negative effect on hippocampal, cerebellar and thalamic volume, a positive effect on lateral ventricular volume, and no effect on putamen volume; baseline age had no effect on cerebellar or putamen volume, a negative effect on hippocampal and thalamic volume, and a positive effect on lateral ventricular volume. Model findings will be reviewed after a summary paragraph highlighting annualized volume change results.

It warrants mention that the use of linear models should not be construed to imply necessarily linear response–predictor relationships. Indeed, in the lateral ventricular linear volume model (based on baseline data, baseline lateral ventricular volume being the response variable) a decidedly nonlinear response–explanatory variable relationship was addressed by a log transform of the response variable ventricular volume (see 4*.1 Linear regression volume analyses*). Nevertheless, regression models were used because of their high interpretability and broad adoption for quantifying variable relationships. In addition, violation of linearity did not occur in models of other structures, nor was there a quadratic shape to the baseline data. In the longitudinal data (just 2-years total time), the mixed model for lateral ventricular volume demonstrated a slight but significantly improved fit by addition of a quadratic term; an indication of some nonlinear change (specifically slight acceleration, expansion) in lateral ventricular volume over time. Finally, it also warrants underlining that, as specified in section 2.5, model results reflect regression not mediation analyses; coefficients quantify the magnitude/strength of the predictor/explanatory/independent variable effect to explain outcome (volume of a given structure). Variables were not assessed as (causal) mediators. For example, in the linear regression analyses thalamic volume as a predictor had consistently the largest coefficient/effect magnitude. This does not validate the thalamus as a causal mediating variable of outcome.

Hippocampal median annualized volume change (see *Supporting information II*, Tables S1-S2), was in agreement with prior research (2, 4, 6, 66), and followed an AD > MCI > CN volume atrophy pattern, with an annual AD hippocampal atrophy of 4.7% that paralleled meta-analytic review findings of 4.66% (^66^). Similarly, the pattern of annualized thalamic volume change (reduction) followed the same atrophy AD > MCI > CN pattern. In annualized normal aging (controls in the current work), and consistent with other research (^67–69^), there was cerebellar, thalamic and particularly hippocampal volume reduction, but lateral ventricular volume increase. In normal aging, however, our lack of change in putamen volume differed from the Fjell et al. (^67^) findings, but was in agreement with another study (^22^). Of relevance, the Fjell et al. research used a much larger sample (*N* = 1100) compared to our project or the Cousins et al., (^22^) study. In addition, in yet other longitudinal research (*N* = 883(^67^)), the putamen also exhibited volume reduction (^69^). It seems likely that the putamen, relative to the other structures, simply requires more data to detect notable volume change. Of note, cerebellum annualized volume change (see section 6.1) was very small (.60 %), an outcome accounted for by slight cerebellar decline in AD as shown in Table 2. This table also suggests that cerebellar volume may be spared in MCI or even greater in MCI. This is in agreement with the consistently higher MCI cerebellar volume that has been reported relative controls (^70^). While AD type pathology may occur in the cerebellum (^71^), it may also be spared in AD (^24^). In short, our findings for cerebellar volume in AD and MCI, like that cited above, are not conclusive. In our mixed model regression analyses (see section *Longitudinal mixed models* that follows the next section) AD pathology (via the group variable) did not contribute to cerebellar volume. But the passage of time significantly contributed to annual reduction in cerebellar volume.

### Linear regression analyses and the primacy of the thalamus

The linear models explaining putamen and cerebellar volume had the highest RMSE and NRMSE values but lowest *adjusted-R**^2^*** values (e.g., putamen volume model, *adjusted-R**^2^*** = .25, RMSE = .05; cerebellar volume model *adjusted-R**^2^*** = .36; RMSE = .70) indicating that the features (explanatory volume variables, age, CDR or group) and algorithm (linear regression) did not amount to a good fit of the observations. By contrast, models explaining hippocampal, thalamic, and lateral ventricular volume had the lowest RMSE and NRMSE but highest *adjusted-R**^2^*** values (e.g., *adjusted-R**^2^***ranged from .52 for a hippocampal volume model to .67 for a thalamic volume model (see **Table 3** for details). Marginally significant interactions occurred in the putamen volume linear model and in one of the thalamic linear models. In both instances age had a moderating but antithetical effect (see *Supporting Information V* for an interpretation).

Importantly, in linear model stepwise (backward using AIC) procedures, thalamic volume, as an explanatory variable, had by far the largest significant estimate in all models (see **Table 3**), as predicted. Thalamic volume had a positive relationship with hippocampal, putamen, and cerebellar volume: a unit increase in thalamic volume explained a (conditional) mean structure volume increase ranging from .38% of total intracranial volume (ICV) in the putamen to 6.3% of ICV in the cerebellum. By contrast, thalamic volume, again as an explanatory variable, had a negative relationship with lateral ventricular volume largely reflecting its anatomical location: the dorsal thalamus forms much of the anatomical floor of the lateral ventricular body (see FIG 1 A). As such, an increase in thalamic volume would encroach on and reduce ventricular volume. Additionally, the hippocampal volume model retained the group variable differentiating AD and MCI pathology cohorts relative to controls. Hippocampal volume mean was significantly reduced in AD and MCI relative to controls (see *4.2* for marginal means and Tukey test results). It was also originally hypothesized that the best fitting thalamic model would also retain the group variable (the proxy for AD and MCI pathology); this hypothesis was not confirmed by the findings.

While the thalamus had the largest linear regression coefficients explaining the volume of other structures, in the model of thalamic volume (thalamic volume as the response variable) the largest coefficients explaining thalamic volume were those from putamen and the hippocampal volume, outcomes in concert with both the critical hub-like role of the thalamus (^34^) and the so-called “rich club network” comprising the thalamus, hippocampus and putamen (^72, 73^).

Moreover, in practical terms, the consistently larger estimates/coefficients of thalamic volume in the role of explanatory/predictor variable positions thalamic volume as the most “bang for the buck” estimate of hippocampal, cerebellar, putamen and lateral ventricular volume. As a single source insight aid of hippocampal, cerebellar, putamen and lateral ventricular volume, the usefulness of thalamic volume is well demonstrated by the current work findings. Moreover, this supports use of thalamic volume in clinical imaging assessments where possible, particularly where the target structure is densely interconnected with the thalamus. Noteworthy, evidence from other research indicates early AD thalamic involvement is confined largely to the anterior thalamic nuclei (^14^). As such, using the entire thalamic volume measure likely diminishes the effect of AD and MCI pathology group membership (^11^). A next step in the current vein of research will include imaging data and analyses derived from isolation of particular thalamic nuclei, such as the anterior thalamic group.

### Longitudinal mixed models

In normal aging, mixed models determined time, passage of just a single year, had a negative effect on hippocampal, cerebellar and thalamic volume, no effect on putamen volume, and a positive effect on lateral ventricular volume. The effect of time on volume, conveyed in FIG 2, FIG 5, **Tables** 4, 5, 6 and annualized volume results, is largely consistent with other research (4, 67-69). As noted earlier in this discussion, the exception was putamen volume. Mixed model longitudinal putamen volume results had the same null change found in the annualized putamen volume outcome. Baseline age had a negative effect on hippocampal and thalamic volume, no effect on cerebellar or putamen volume and a positive effect on lateral ventricular volume. With the exception of the putamen, these findings are in keeping with other research (^12^) employing the same segmentation platform. Over the adult life span, a smoothing spline model captured accelerated change for the lateral ventricles, cerebellum, putamen and particularly the hippocampus (^67^). Similarly, over the course of a decade, non-linear patterns of hippocampal and lateral ventricular volume change have been reported (^69^). Over the course of a single year, accelerated hippocampal atrophy measured by quadratic expansion (6), has been reported. The present work mixed model findings included a small but significant quadratic time term indicative of annual marginally accelerated lateral ventricular volume increase (see **Table 6**). Specifically, the quadratic term for time was positive (as was the effect of time itself, see **Table 6**) indicating a significant incremental effect of time on lateral ventricle volume (*p* = .03). This is consistent with the FIG 5 (B) lateral ventricular volume plot, where a slight convexity appears (in the mean thicker loess lines).

The best fitting hippocampal and thalamic mixed models retained the group variable, the proxy of AD or MCI potential AD pathology. The marginal (group) predicted hippocampal volume means, at baseline age, and for all three time points were significantly (*p* < .05) lower for both AD and MCI relative to controls. Group predicted hippocampal AD and MCI mean volumes did not significantly differ. Cohen’s *D* values (AD, *d* = -1.9; MCI, *d* = -1.56) indicated large negative effects of AD and MCI group membership on hippocampal volume relative to controls.

Thalamic marginal means were also lower in AD and MCI groups relative to controls, but only the MCI marginal mean significantly differed (z = -2.58, *p* = .03) from controls. While thalamic AD vs control predicted marginal means did not significantly differ (z = -2.12, *p* = .09, *d* = -.74) the Cohen’s *d* effect sized of -.74 indicates a moderate strength difference between AD and control group thalamic volume. See *6.2 Mixed model results* for details.

Large negative AD and MCI Cohen’s *d* effect sizes for the hippocampus (sections *4.2* and *6.3*) indicate proportionally large, negative effects of AD and MCI membership on hippocampal volume relative to controls. The strength of group membership was not as pronounced for thalamic volume. This may, at least in part, be due to use of the entire thalamus volume measure, which, as noted at the end of the section *Linear regression analyses and the primacy of the thalamus,* likely diminishes model sensitivity. More pronounced hippocampal longitudinal volume decline in AD has been previously reported (4, 6, 66, 74), as has greater thalamic volume decline in AD (4). Retention of the group variable in both the hippocampal and thalamic models, in the present work, is in keeping with the well documented interconnectedness of these structures (^33, 38, 39^), evidence of early AD pathology in both structures (^14, 75^), and also consistent with repeated measure correlation findings (see **Table 4**), where the second strongest correlation (*rrm* (71) = .74, *p* < .0001) occurred between hippocampal and thalamic volume. Both AD and MCI thalamic volume have exhibited reduction in other research (4, 15, 16).

The present work findings, are in line with research from Braak et al. (^13, 14^) and Aggleton et al. (^11^) indicating early-stage AD type pathology in both hippocampus and thalamus. The mixed model group variable (again representing AD or MCI pathology relative to controls) was not retained by best fitting/parsimonious cerebellar, putamen and lateral ventricular volume mixed models but was retained by the hippocampal and thalamic models (See FIG 5 and **Tables 5, 6**). This outcome in the early-stage cohorts assessed in the present work is consistent Braak et al. and Aggelton et al findings.

Of note, at the time of this writing, a PubMed search (e.g., ((Alzheimer’s) in Abstract) and ((hippocampal volume) in Abstract) and ((mixed model) in Abstract)) returned several studies utilizing mixed models in Alzheimer’s disease research. However, none conducted univariate mixed models using volume of the subcortical structures (thalamus, cerebellum, lateral ventricles, putamen) employed in the current work as the dependent variables. Some studies did include hippocampal volumetry but it seems only one longitudinal study (^76^) used hippocampal volume as the response variable and also provided a normalized mixed model estimate (fixed effect coefficient) for the annual effect of time on hippocampal volume. The latter referenced study focused on MCI and used a metric of normalized volume in mm^3^. When the mm^3^ estimate was converted to percentage of total intracranial volume (ICV, the volume measure used in the current work) the mixed model coefficient for time (as the solitary predictor or included with other predictors, such as age) was in a range similar to the current work (-.01 in our model; -.02 to -.03 in the Huijbers et al (^76^) study).

In summary, thalamic volume linear regression explanatory estimates made the greatest contribution to variation in hippocampal, cerebellar, putamen and lateral ventricular volume.

Mixed models determined the passage of a single year increased lateral ventricular volume, reduced hippocampal, cerebellar and thalamic volume, but had a negligible effect on putamen volume. Baseline age had a negative effect on hippocampal and thalamic volume, no effect on cerebellar or putamen volume and a positive effect on lateral ventricular volume. Moreover, the group (proxy for AD or potential pathology in MCI) variable’s contribution to significant volume variation was confined to hippocampal volume in linear regression. But in mixed longitudinal models the group variable significantly contributed to both the hippocampal and thalamic volume variation. This is one indication that linear models were not as effective as linear mixed models discriminating early AD pathology in the thalamus. Repeated measure models generally have more statistical power (^77^). It is assumed this enabled the mixed model to detect early AD pathology not just in the hippocampus but in the thalamus as well.

Thalamic volume, the pivotal linear regression imaging-derived predictor in the present work, may, speculatively, along with specific thalamic nuclei(^11^) metrics, provide centralized insight to AD intervention effects on brain structures with dense thalamic interconnections.

There is a growing tide of evidence, linking fasting (such as intermittent fasting) (^78–83^) and exercise interventions to augmented neurogenesis (^84–88^) - neurogenesis is the antithesis of the volume reduction correlate neurodegeneration. Supervised exercise and intermittent fasting offer immediate, substantiated therapeutic and potentially prophylactic benefits for AD (^84–88^). In the near future, induced pluripotent stem cell (iPSC) technology (^89–91^) will likely be at the forefront of AD and other pathology clinical interventions. For a mini up to date review of exercise, intermittent fasting, iPSC technology and other research relevant to treatment of AD see Mitigating Neurodegeneration in *Supplementary Information III*. Where neuroimaging data is available, the primacy of the thalamus as a predictor in the current work supports thalamic metrics, volume validated here, to aid in intervention assessments.

Undoubtedly, the understanding of AD will be advanced by research incorporating intermittent fasting, exercise and eventually iPSC technology interventions. Imaging will continue to play a vital role, non-invasively revealing the effects of such interventions. Finally, mixed models offer standard regression quantification of intervention effects over time. Broader adoption of mixed models would promote much improved method consistency across longitudinal study analyses.

## Limitations

Relatively small sample size (*N* =45 in linear models; *N* of 33-36 in mixed models) and group size (minimum *n* = 10 AD in the thalamic mixed model) raise understandable concern about reproducibility and generalization. In general, an *N* ≥ 30, according to the central limit theorem, approximates the population mean and variance. A minimum of *N* ≥ 25 has recently been estimated for regression models (^92^). With regard to group *n,* a publishing minimum requirement of *n* = 5 has been stipulated (^93^). As specified above, this work complied with these *N* and group *n* minimums, suggesting plausible sample to population model inference. It must also be cautioned that results could have been impacted by selection bias: data was not randomly selected but filtered by age, scan resolution and interpreted scan quality. In addition, gender imbalance (notable in the AD cohort comprised of 27 females and just 3 males) was an unintended consequence of the data filtering. The filtering may have inadvertently selected individuals with less genetic-based volume differences, which in turn would diminish a distinguishing gender variable effect. Finally, evidence of thalamic pathology in early AD is localized largely to the anterior thalamic nuclei. Consequently, use of only whole thalamic volume likely diminished thalamic volume coefficient/effect size. Nevertheless, evidence of detectable whole thalamic volume alteration in AD and in normal ageing has been reported elsewhere (4, 12, 67). Most clinical imaging assessments use whole structure volumes. For these reasons, whole thalamic volume was used in the current work.

## Conclusion

Volume, particularly hippocampal volume, is a widely researched metric in AD. Status of other structures in AD, such as the cerebellum, putamen, lateral ventricles and thalamus, have collectively undergone less scrutiny. Current work linear regression analyses underlined the utility of thalamic volume as a virtual centralized index of hippocampal, putamen, cerebellar and lateral ventricular volume. This is in agreement with but extends introduction literature findings by further validating the seemingly pervasive influence of the thalamus. Thalamic volume lent unequalled single-source insight to volume in the other structures and is well supported in the current work as an explanatory variable of high potential, particularly where the target structure is densely interconnected with the thalamus. In addition, mixed longitudinal models discriminated and quantified evidence of early AD pathology not just in the hippocampus but also in the thalamus. This too is congruent with introduction literature findings reporting early AD stage pathology in these two structures, and supports including thalamic in addition to hippocampal volume in early-stage AD assessments. The current study findings justify additional similar research using larger samples. The next phase of this research, with similar but broadened scope, will include assessment of volume and connectivity relationships among particular thalamic nuclei (e.g., anterior thalamic nuclei) and other structures, larger samples and a cross-validated approach.

## Disclosure statement

The authors stipulate there were no financial or other relevant influential interests potentially biasing research. This research did not receive funding from any specific non-profit or commercial agency.

## Conflict of interest

The authors stipulate there was no conflict of interest.

## Author Contributions

Verification of CSL^†^’s analyses conducted by MH. Overall proofing and edit suggestions conducted by WDS and JFD. First authorship = ^†^.

## Data availability statement

Data used is available directly from ADNI’s data access repository (adni.loni.usc.edu/data- samples/access-data/). The specific sample used can also available from GitHub (https://github.com/csl4r/Subcort_Volume) and this website (supporting information Datasets).

The R code can be obtained from the first author anytime.

## Abbreviations

ADNI = Alzheimer’s Disease Neuroimaging Initiative; AD = Alzheimer’s disease; CN = controls; MCI = mild cognitive impairment; BTB = MRI T1 same-subject scans taken in back-to -back; CDR = clinical dementia rating; MMSE = mini mental state exam; MPRAGE T1 = Magnetization Prepared Rapid Acquisition Gradient EchoT1 weighted image; NRMSE = normalized root mean squared error; V0A and V0B = baseline BTB same-subject scan pairs; V12A and V12B = 12-month BTB same-subject scan pairs.

## Notes

### Competing Interest Statement

The authors have declared no competing interest.

